# Natural and synthetic potential inhibitors of endoribonuclease Nsp15 encoded by Middle East Respiratory Syndrome Coronavirus (MERS-CoV). Computer modeling *in silico* experiments. *p*-Coumaric acid, Curcumin and their Boronic acid derivatives compared to Hydroxychloroquine and its Boronic acid derivative

**DOI:** 10.1101/2020.03.23.002881

**Authors:** Juan F. Barquero

## Abstract

Based on the structural and biochemical characterization of endoribonuclease Nsp15 in crystal structure PDB code 5YVD, I am providing plausible inhibitors of this enzyme. In this report I intent to signal that is possible to inhibit this enzyme by the use of natural occurring compounds and their boronic acid derivatives, compounds with borono B(OH)_2_ groups. Boronic acids are atracted to serine side chains in the active site of serine proteases, in this case an endoribonuclease. Actual lab tests need to be conducted, nevertheless I venture here to propose four compounds: *p*-coumaric acid, Curcumin and their boronic acid derivatives with the use of computer modeling, *in silico* experiments. I also compared the above mentioned compounds to Hydroxychloroquine and its boronic acid derivative. I used *AutoDock Vina, UCSF Chimera* 1.12, *DeepView / Swiss-Pdb Viewer* 4 1.0, *Sulp* 3.0, *PubChem, MDL Isis Draw* 2.5, *OpenBabelGUI* 2.2.3, and *Microsoft WordPad*. As hardware I used a VAIO loptop with Intel Core i5 with *Microsoft Windows* 8.0, and a Dell loptop Inspiron E 1405 with an Intel Core Duo with *Microsoft Windows* Xp (off line).

## Introduction

The crystal structure with PDB code 5YVD corresponds to Nonstructural protein 15 (Nsp15) that is encoded by coronavirus (CoV). This protein is crucial in the life cylcle of coronavirus (CoV) responsible for Middle East respiratory syndrome CoV (MERS-CoV). This protein may be ubiquitous to corona viruses, the Nsp15 of MERS-CoV, severe acute respiratory syndrome coronavirus (SARS-CoV), and mouse hepatitis virus (MHV) share homology. Moreover, Nsp15 bind to other proteins such as Nsp8 and Nsp7/Nsp8 complex that are involved in replication and transcription of RNA. If one of the enzymes in the metabolic cycle of the virus is inhibited then the spread of the virus may be halted. Moreover, according to Zhang *et al*., when Ser290 of chain A of Nsp15 is mutated, substituted by alanine, the activity of enzyme is reduced by 67% of the wild type Nsp15. Ser290 confers uridylate specificity to the enzyme. (Zhang *et al*., 2018). Therefore, we have an important enzyme in the coronavirus cycle, we have an active site, and we have a specific target in the active site - Ser290. On the other hand, we know that boronic acid target Serine side chains in serine proteases, we just have to test them against this particular endoribonuclease.

*p*-coumaric acid occurs in the plant world in abundance and it shows biological activity. (Hoffmann, 2003). Other names: Phloretic acid, desaminotyrosine. IUPAC name: 3-(4-hydroxyphenyl)propanoic acid. Also an IUPAC name: 3-(4-hydroxyphenyl) propanoic acid. CID in PubChem: 63542 (PubChem).

Curcumin shows antioxidant and anti-inflamatory activity, it is found in tropical spices. (Hoffmann, 2003). Other names are: Natural yellow 3, tumeric yellow, tumeric, curcuma, Indian saffron, diferuloylmethane, and the IUPAC name (1E, 6E)-1,7-bis(4-hydroxy-3-methoxyphenyl)hepta-1,6-diene-3,5-dione. The CID: 965516 in *PubChem*. (*PubChem*).

Hydroxychloroquine, with IUPAC name: 2-[4-[(7-chloroquinolin-4-yl)amino]pentyl-ethylamino]ethanol. (PubChem), An antimalarial drug, controversial for its presumptive toxicity, nevertheless used in this report to compare to the other compounds.

Boronic acids are the key to attack a specific site in the active site of this enzyme, and that specific site is serine 290 in the case of chain A of the Nsp15 hexamer. Boron is an atom that is electron deficient, it contains an electron sink, where atoms that are electron rich can deposit the extra electrons. This makes the boron a Lewis acid and not a Brønsted-Lowry acid (Bruice, 1995). Boronic acids show reversible competitive inhibition. Boron form a tetrahedral transition-state with the serine of the active site also because it easily interconverts between sp^2^ and sp^3^ hybridization. (DeSoyza, 1990) (Weston, 1998) (Hall, 2005). (See figures 1 and 2).

**Figure 1:**
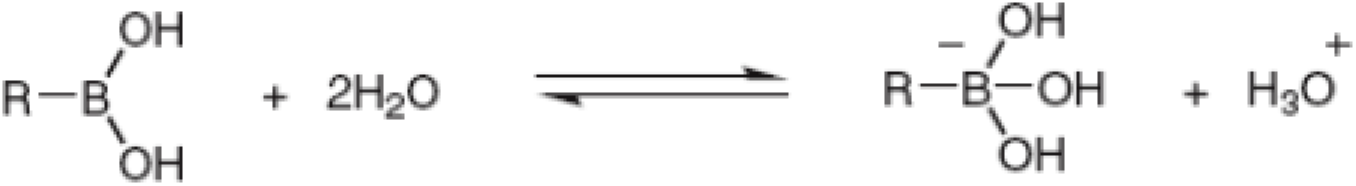
Ionic equilibrium of boronic acid in water, sp^2^ and sp^3^ hybridization equilibrium. (From Hall, 2005).

**Figure 2.**
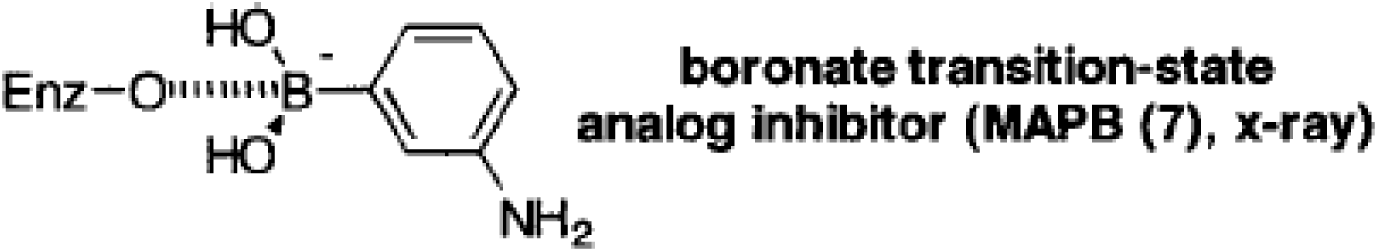
Boronic acids as transition state analogues, sp^3^ hybridization (From Weston *et al*., 1998). *m*-aminophenylboronic acid (MAPB) (Weston *et al*,. 1998) (Steinberg, 1964) (Usher, 1998).

Boronic acids have low toxicity compared to other organic compounds, they show little threat to the environment (Hall, 2005) (Tafi, 2005).

**For ease of communication where I added a boron to the structure I added a B for boron at the end of the common name; *p*-coumaric acid becomes *p*-coumaricB acid, Curcumin becomes CurcuminB, and Hydroxychloroquine becomes HydroxychloroquineB**.

It is important to predict the binding of substrate to enzyme in an efficient manner before conducting an actual experiment in the lab (Oleg *et al*., 2009). I used as a predictive instrument the program *AutoDock Vina*. As a docking program, *Vina* provides a score based on energy minimums Gibbs free energy (ΔG) or an association constant given by the relation ΔG = -RTlnK, where K is an association constant. Each conformation of a ligand in the active site presents a ΔG in *Vina*. The lower the ΔG the more favorable. Because K is an association constant then the reciprocal represents a dissociation constant 1/K called ki which represents the inhibition constant for that ligand (See derivation below).

*UCSF Chimera* 1.12 interfaces with *AutoDock Vina*, and provides the first glance at the results of the docking procedure.

*Sculpt* 3.0 is used for converting the docked ligand into pdb file for use with *Sculpt* 3.0 and *Swiss-PdbViewer* 4 1.0 pdb’s by *UCSF* Chimera don’t work well in *Sculpt* and *Pdb Viewer. Sculpt* lets visualize the active site determined by a surface at a distance of 10 Angstroms from the ligand. It is important to know which amino acid side chains are in the active site.

*Swiss-PdbViewer* 4 1.0 is used to merge the ligand and the receptor enzyme into one pdb file for processing after the docking has been executed by *Vina*. It is also used for visualization of ligand-enzyme distances and hydrogen bond lengths.

*MDL Isis Draw* 2.5 was used to draw the molecules of boronic acids in 2D.

*OpenBabelGUI* 2.2.3 is used to transform *Isis Draw* files to pdb files with 3D coordinates.

*PubChem* (on line) is used to draw the natural occuring molecules.

*Microsoft WordPad* is of crucial importance in numbering the ligand molecule in the newly merged receptor-ligand by *Swiss-Pdb Viewer*. The number assigned is the next number in the amino acid sequence, in this case 342. Once this number is assigned in the pdb file, the active site can be found in *Sculpt* and distances receptor-ligand in *Swiss-Pdb Viewer*.

## Procedures

### Preparation of the enzyme

From the National Center for Biotechnology Information (NCBI) website, a Protein Data Bank crystal structure of Nsp15 (PDB code: 5YVD) was downloaded.

Chain A of the hexamer 5YVD was isolated using *UCSF Chimera* 1.12.

A glycerol molecule present in the active site of 5YVD was removed.

### Preparation of the ligand

The naturally occuring compounds *p*-coumaric acid, and curcumin as well as glycerol were drawn in *PubChem* and saved as pbd files. These structures were then minimized using *UCSF Chimera* 1.12 (Dunbrack, 2002).

Boronic acid structures were drawn in *MDL Isis Draw* 2.5, and then converted to pdb files with *OpenBabelGUI* 2.2.3. Then minimize with *UCSF Chimera* 1.12.

### Conducting the *in silico* experiment (Docking)

Both the receptor enzyme and the ligand were opened in *UCSF Chimera* 1.12. The pre-dock procedure (Dunbrack, 2002) was conducted from the tools menu, and then the actual docking by *Vina’s* interface.

In *AutoDock Vina* we need to specify a center for a box that engolfs the active site. I made it the center the coordinates of carbon number 2 of glycerol C_2_, the original glycerol that shows in the 5YVD chain A crystal structure. The parameters are: 86.487, 154.530, and 40.433. The box was made 20, 20 and 20 units in dimention.

The results were recorded as pdbqt files. These pdbqt files were saves as text in *Word pad* as pdb’s. All six outcomes are one after the other in this pdb file. The best model was selected and then saved again on a separate file.

The pdb of the ligand was transformed into pdb written by *Sculpt* 3.0 by open it in *Sculpt* and then saving it as pdb. This file was then unified to the pdb of the receptor enzyme (5YVD) in *Swiss-PdbViewer* 4 1.0 by saving (project) all layers.

Now the file of both the ligand and the receptor unified by *PdbViewer* is opened by *Sculpt* and saved as pdb file again. Then it was opened with *WordPad* to change the number of the ligand to the next number in the amino acid sequence of the receptor, number 342.

### Viualizing enzyme-ligand interaction

Now the file can be easily visualized In *Sculpt* and *PdbViewer*. In *Sculpt* I found out the distance from ligand to the backbone of the amino acids from 5 to 10 and some cases 12 Angstroms away. And in *PdbViewer* I found Hydrogen bonds and their distances to the receptor.

Note that because my version of *Vina* doesn’t recognize boron, carbon is not substituted for boron in this experiment, the only added advantage is that boronic acid would have an extra hydrocyl group increasing the atraction to Ser290 by means of hydrogen bonds. So, both structures *p*-coumaric acid and its boronic acid derivative would be the same. In the case of Curcumin and CurcuminB, they are not the same and therefore both structures were docked.

The figures in the results section of this report are the ones in which the ligand is oriented or closer to Ser290 to prove the point that Ser290 should be our target in the active site.

### Computation of inhibition constant (ki) from ΔG (affinity constant) from *AutoDock Vina*

ΔG = -RTlnK,

Where K is an association constant, R = 8.3144 J K^-1^ mol^-1^, T is temperature in Kelvin (Eisenberg, 1979), (Martin, 2020).

K = e^-(ΔG/RT)^

ki = 1/K, where ki is a dissociation constant ki = 1/ e^-(ΔG/RT)^

ki = e^(ΔG/RT)^

ki x 10^6^ = ki in the micro molar (µM). (University of Oulu website).

Vina’s ΔG is temperature independent. Assumed biochemistry standard temperature of 25 °C (298.15 K). R in kcal/mol = 0.001987 kcal/mol

RT for our purposes is then 0.592451 kcal/mol

Ki = (e^(ΔG/0.592451)^) x 10^6^ Where ΔG is what is reported by *Vina*.

Result = Ki in micromolar. We recognize units of micro molar even though the computations end unitless because moles were imbedded in the computations.

**Figure 3.**
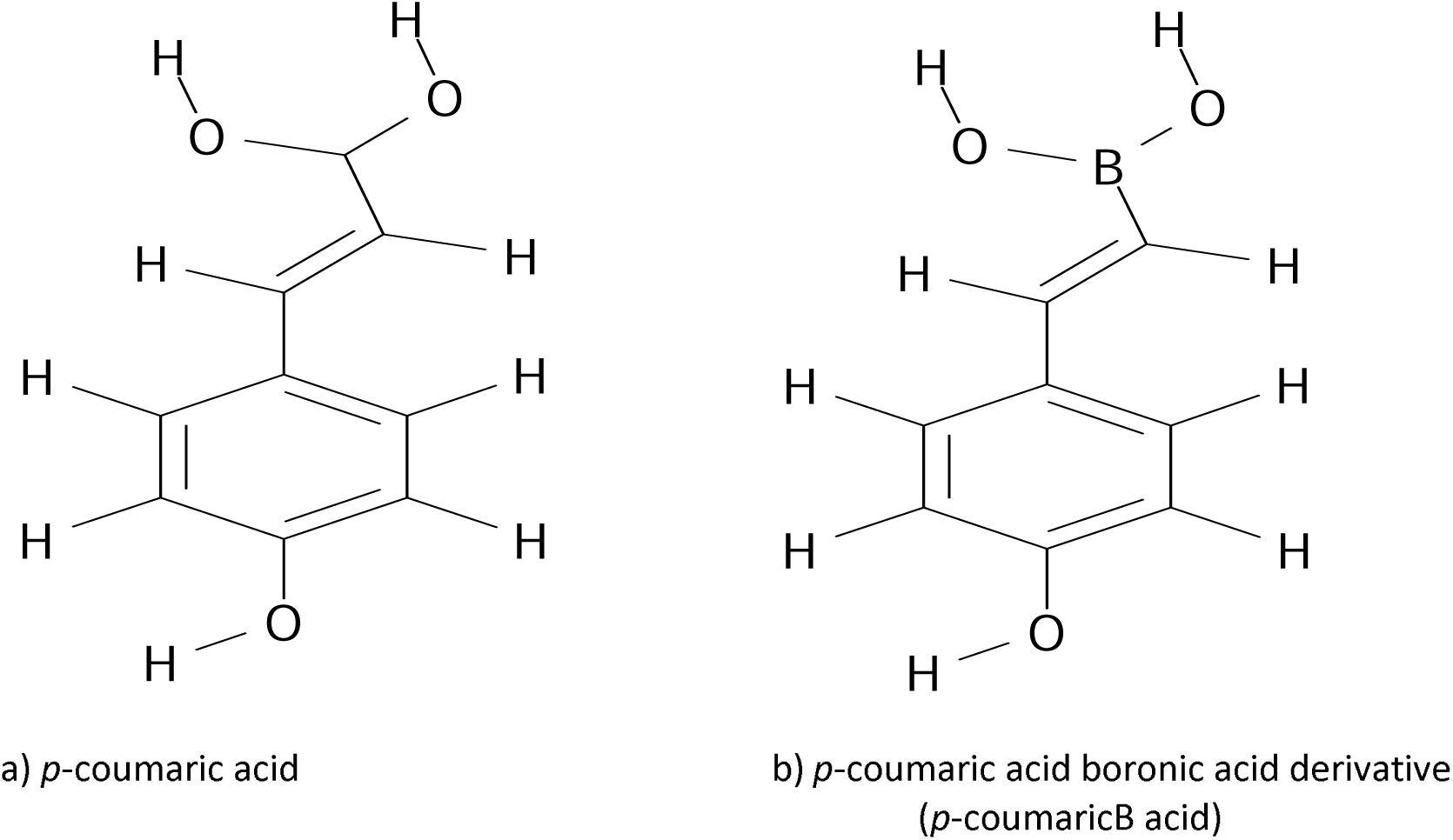
Side by side comparison of *p*-coumaric acid and its boronic acid derivative. Drawn with *MDL ISIS Draw* 2.5. (*MDL ISIS Draw* 2.5).

**Figure 4.**
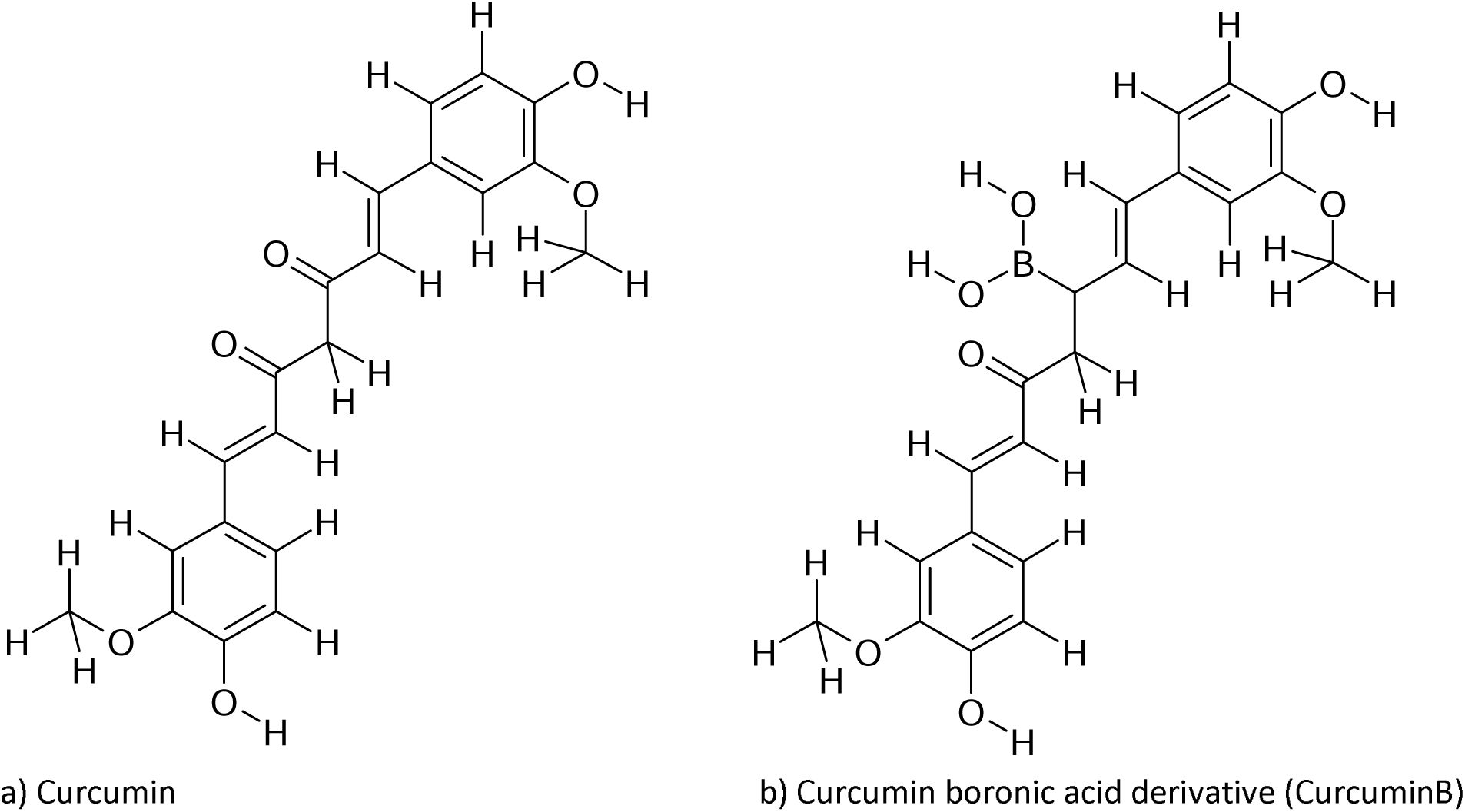
Side by side comparison of curcumin and its boronic acid derivative. Drawn with *MDL ISIS Draw* 2.5. (*MDL ISIS Draw* 2.5).

**Figure 5.**
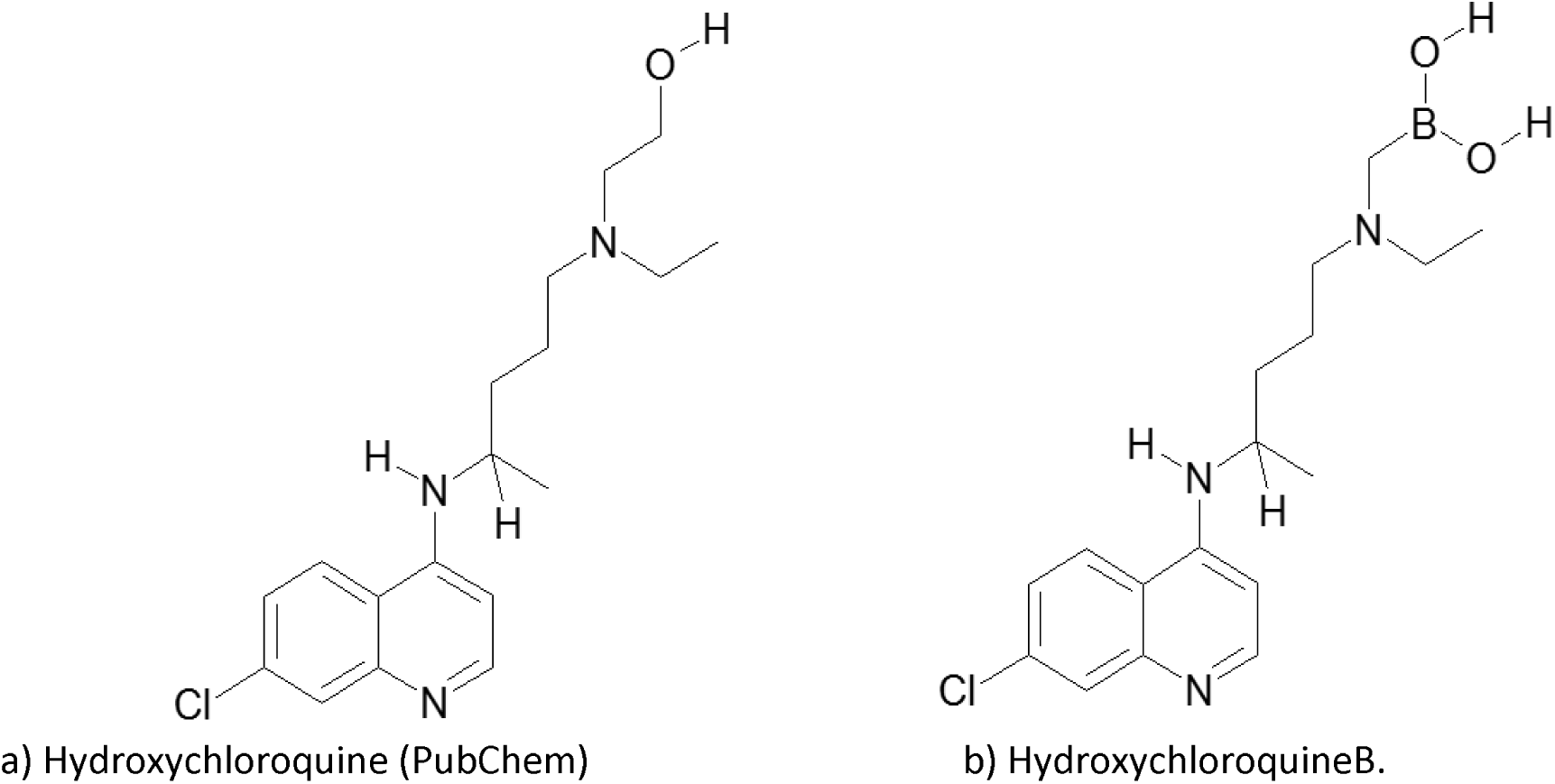
Side by side comparison of Hydroxychloroquine and its boronic acid derivative HydroxychloroquineB. *MDL ISIS Draw* 2.5.

## Results

**Table 1.**
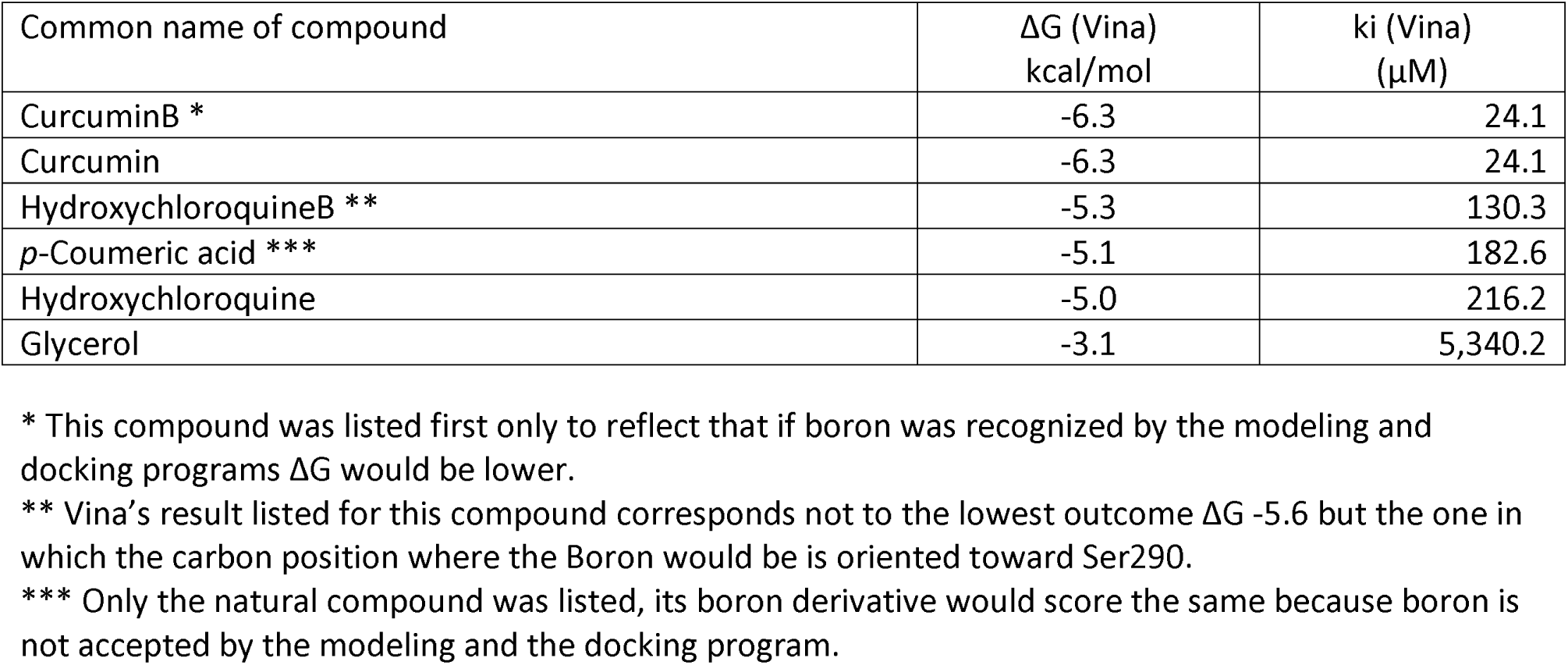
Free energy per Vina’s results converted to inhibition constant (ki) in micromolar.

| Common name of compound | $\Delta G$ (Vina)<br>kcal/mol | $k_i$ (Vina)<br>($\mu M$) |
| --- | --- | --- |
| CurcuminB * | -6.3 | 24.1 |
| Curcumin | -6.3 | 24.1 |
| HydroxychloroquineB ** | -5.3 | 130.3 |
| *p*-Coumeric acid *** | -5.1 | 182.6 |
| Hydroxychloroquine | -5.0 | 216.2 |
| Glycerol | -3.1 | 5,340.2 |

\* This compound was listed first only to reflect that if boron was recognized by the modeling and docking programs $\Delta G$ would be lower.
\*\* Vina's result listed for this compound corresponds not to the lowest outcome $\Delta G$ -5.6 but the one in which the carbon position where the Boron would be is oriented toward Ser290.
\*\*\* Only the natural compound was listed, its boron derivative would score the same because boron is not accepted by the modeling and the docking program.

### Results for *p*-coumeric acid

**Figure 6.**
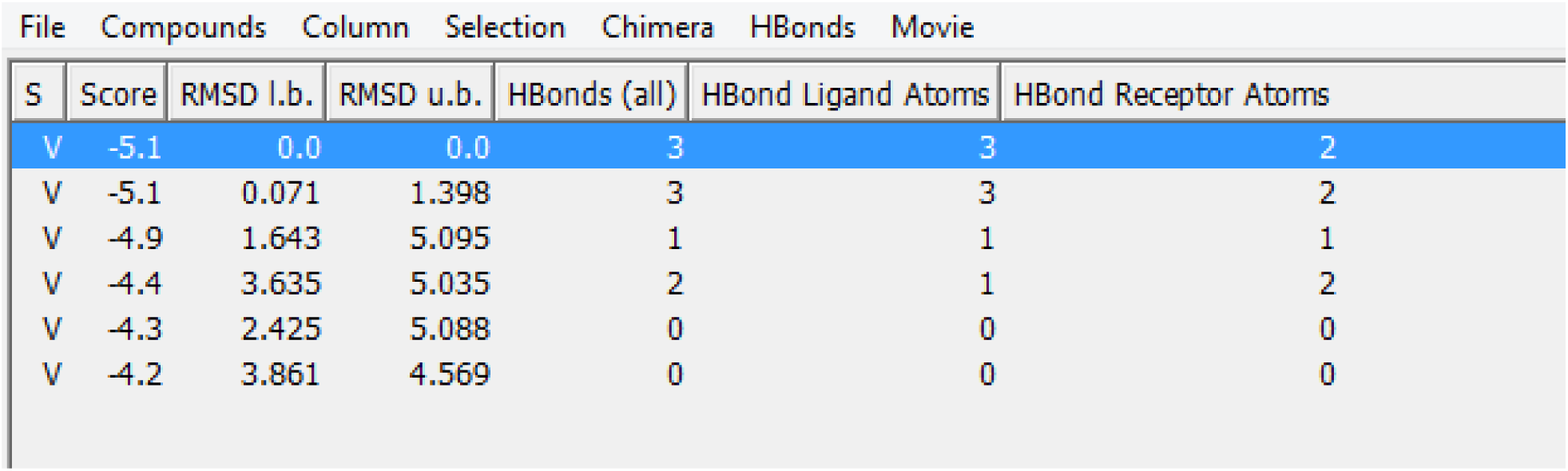
*AutoDock Vina* results. Docking of *p*-coumeric acid and Nsp15 (5YVD A). *UCSF Chimera* 1.12.

**Figure 7.**
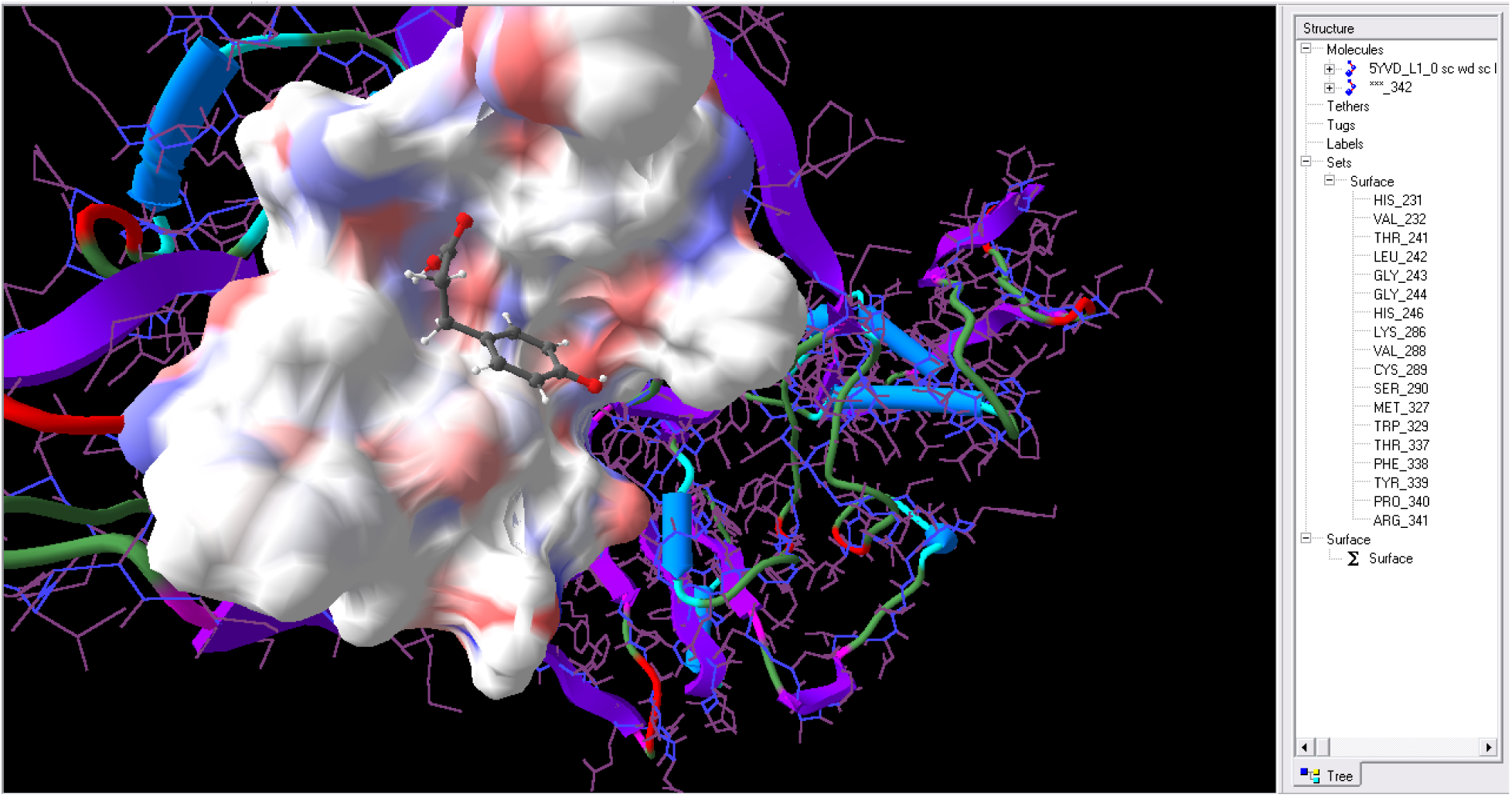
*p*-coumaric acid in the active site of Nsp15 (5YVD A). *Sculpt* 3.0 List of amino acids on the right side are within 10 Angstroms of the ligand.

**Figure 8.**
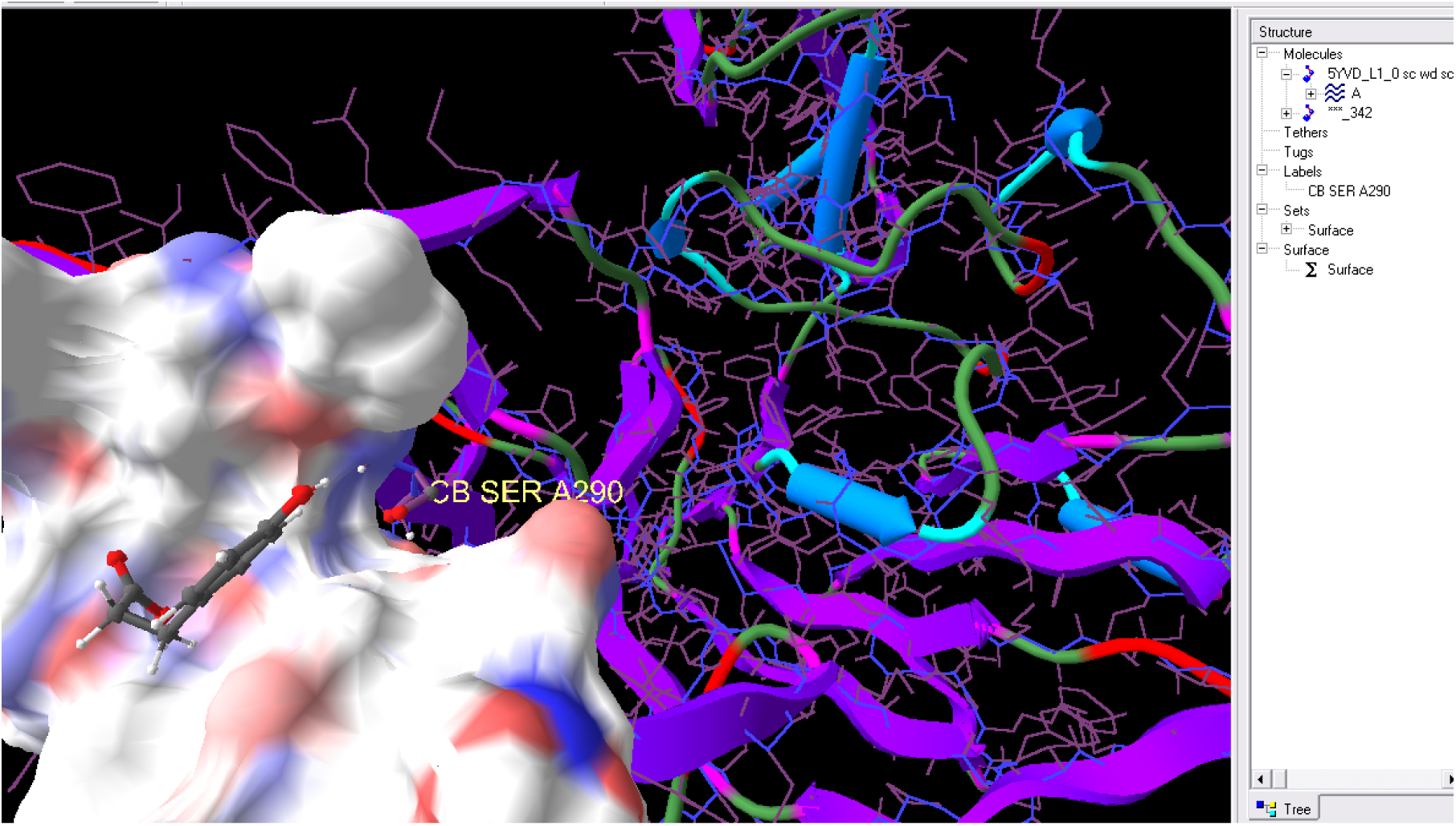
*p*-coumaric acid in the active site of Nsp15 (5YVD A). *Sculpt* 3.0 The surface was removed from Ser290 which is part of the active site. Ser290 appears at 8.00 Angstroms from the ligand.

**Figure 9.**
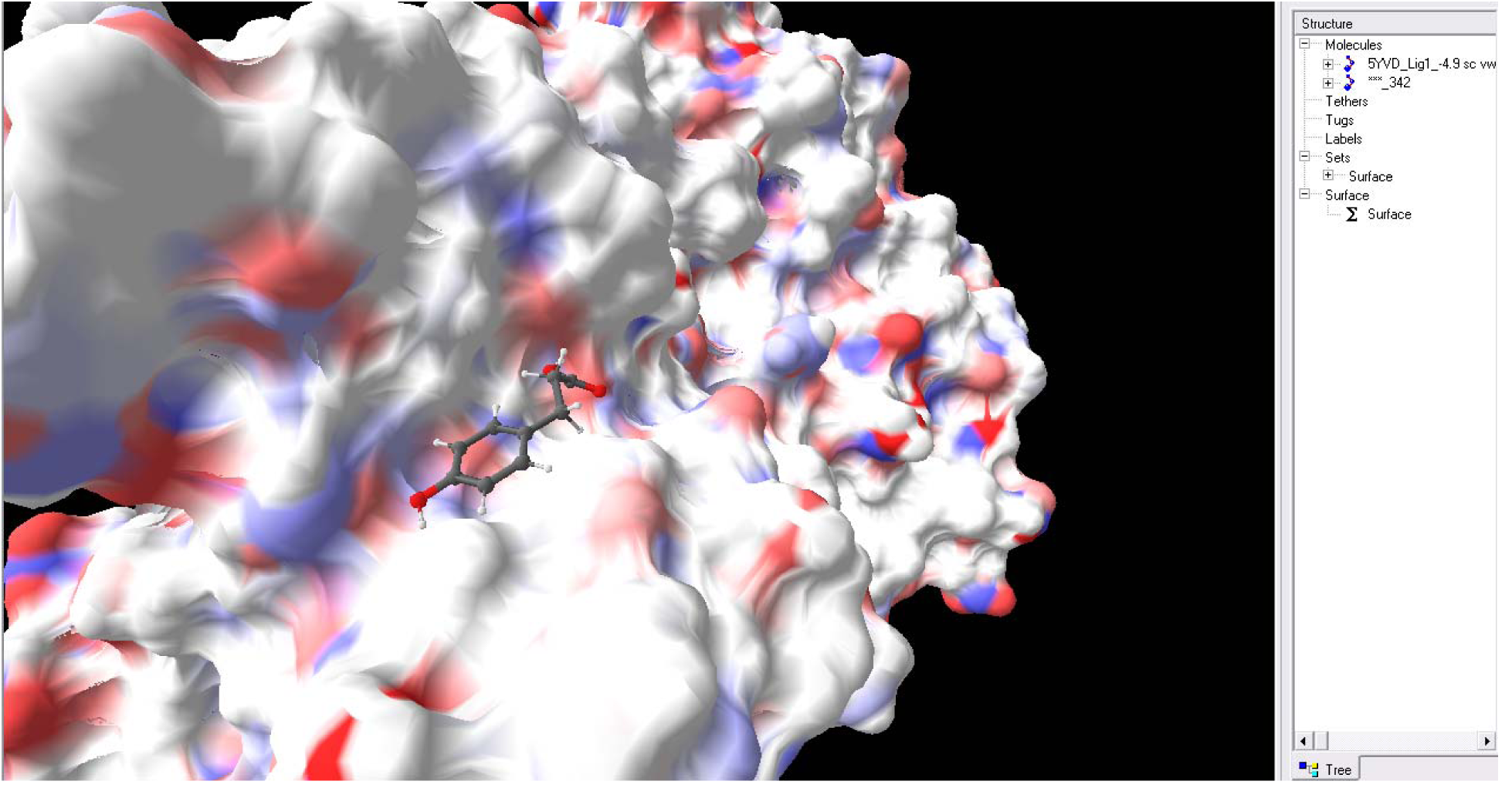
*p*-coumaric acid in the active site of Nsp15 (5YVD A). *Vina’s* free energy −4.9. Note proximity of ligand to Ser290 *Sculpt* 3.0.

**Figure 10.**
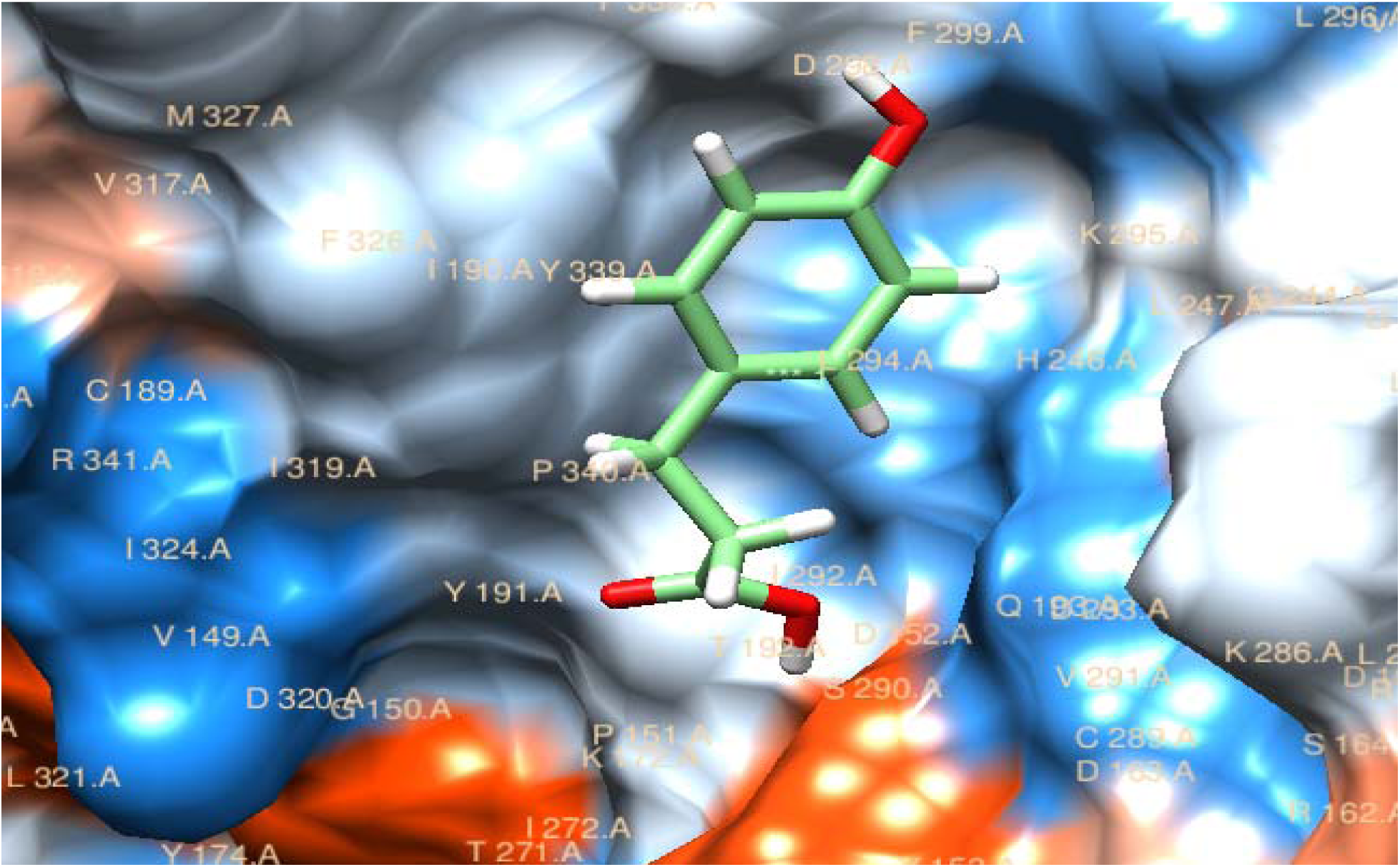
*p-*coumaric acid in Nsp15 (5YVD A). *AutoDock Vina*’s free energy −4.9. Note proximity of ligand to Ser290. *UCSF Chimera* 1.12

**Figure 11.**
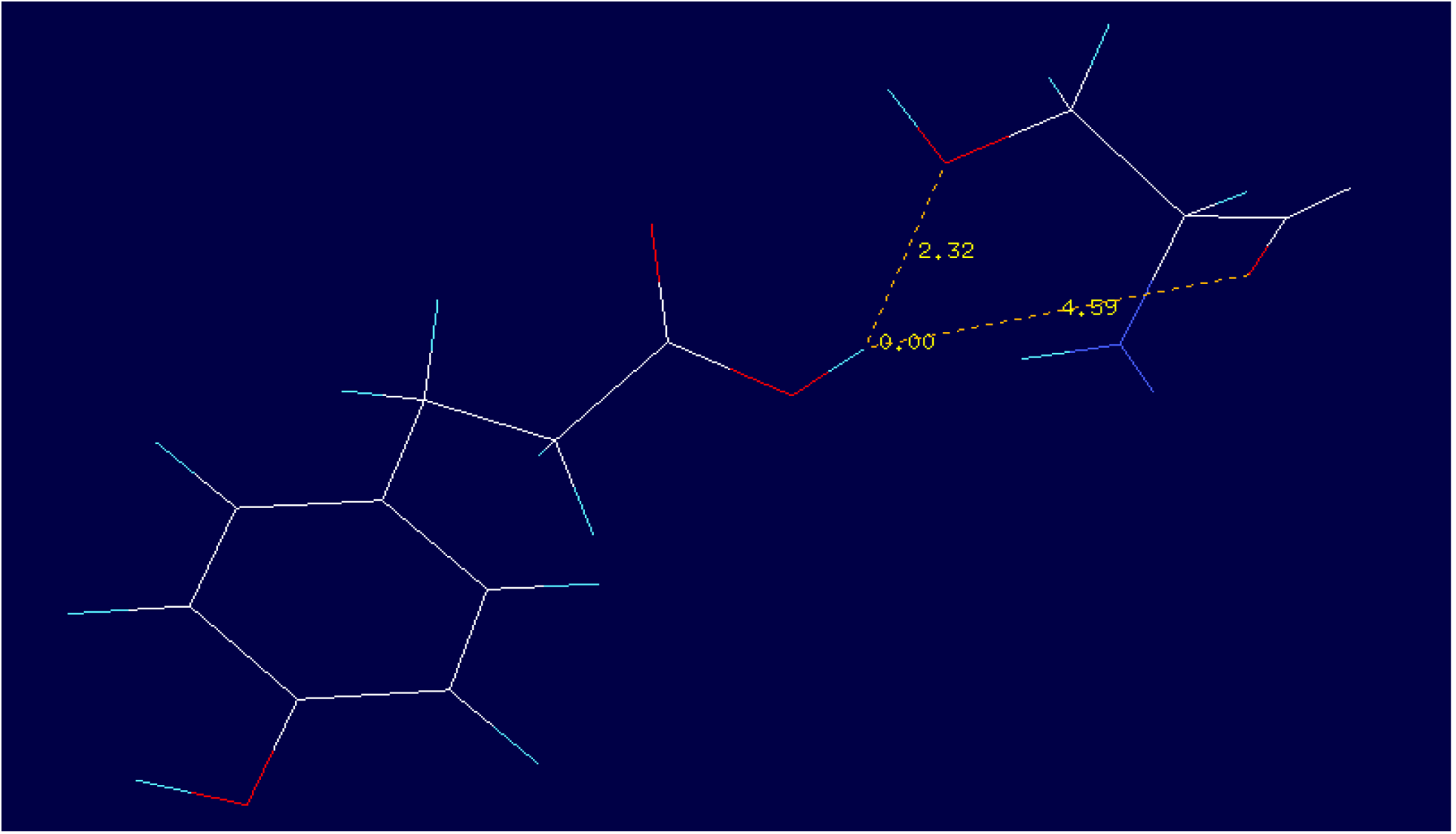
*p-*coumaric acid in Nsp15 (5YVD). *AutoDock Vina*’s energy = −4.9. *UCSF Chimera* 1.12

### Results for Curcumin

**Figure 12.**
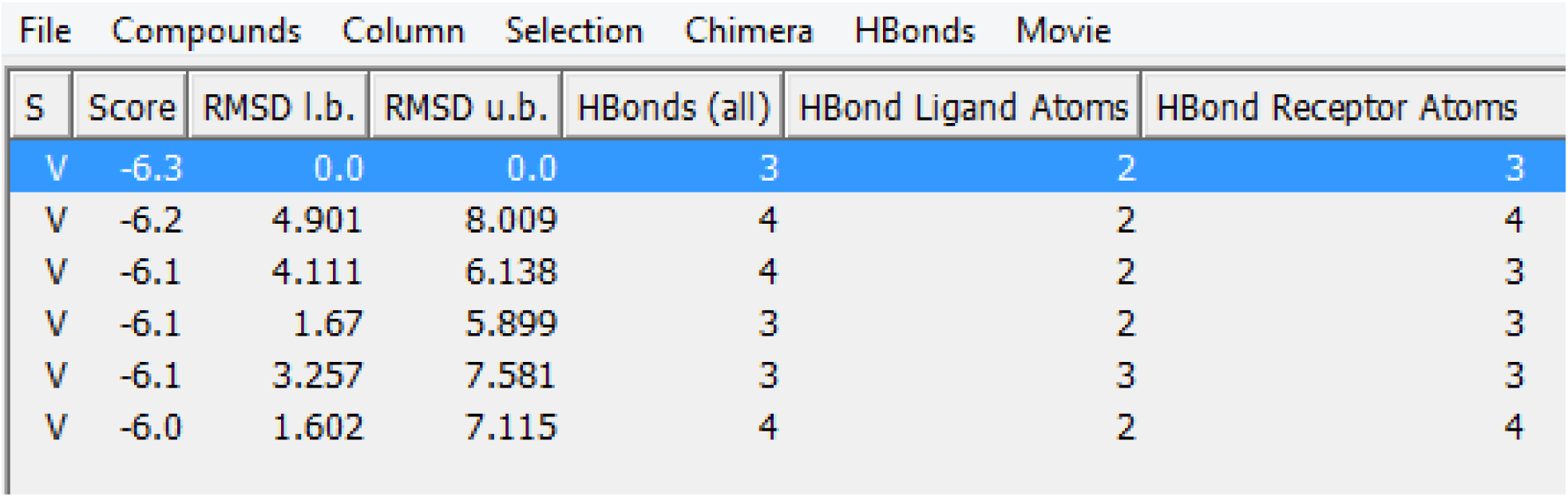
*AutoDock Vina*’s docking of Curcumin and Nsp15 (5YVD A). *UCSF Chimera* 1.12.

**Figure 13.**
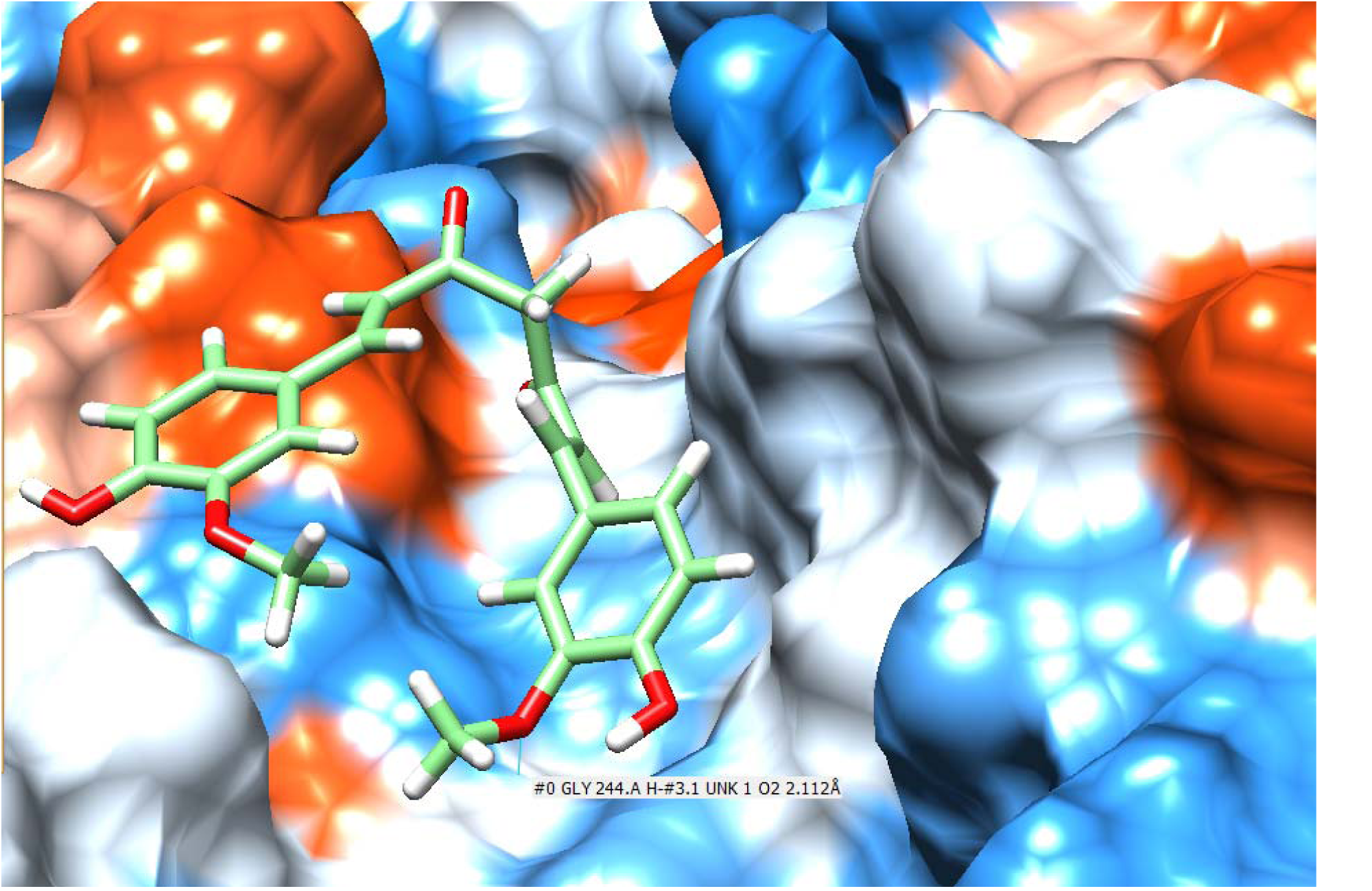
Curcumin in active site of Nsp15 (5YVD A). *UCSF Chimera* 1.12.

**Figure 14.**
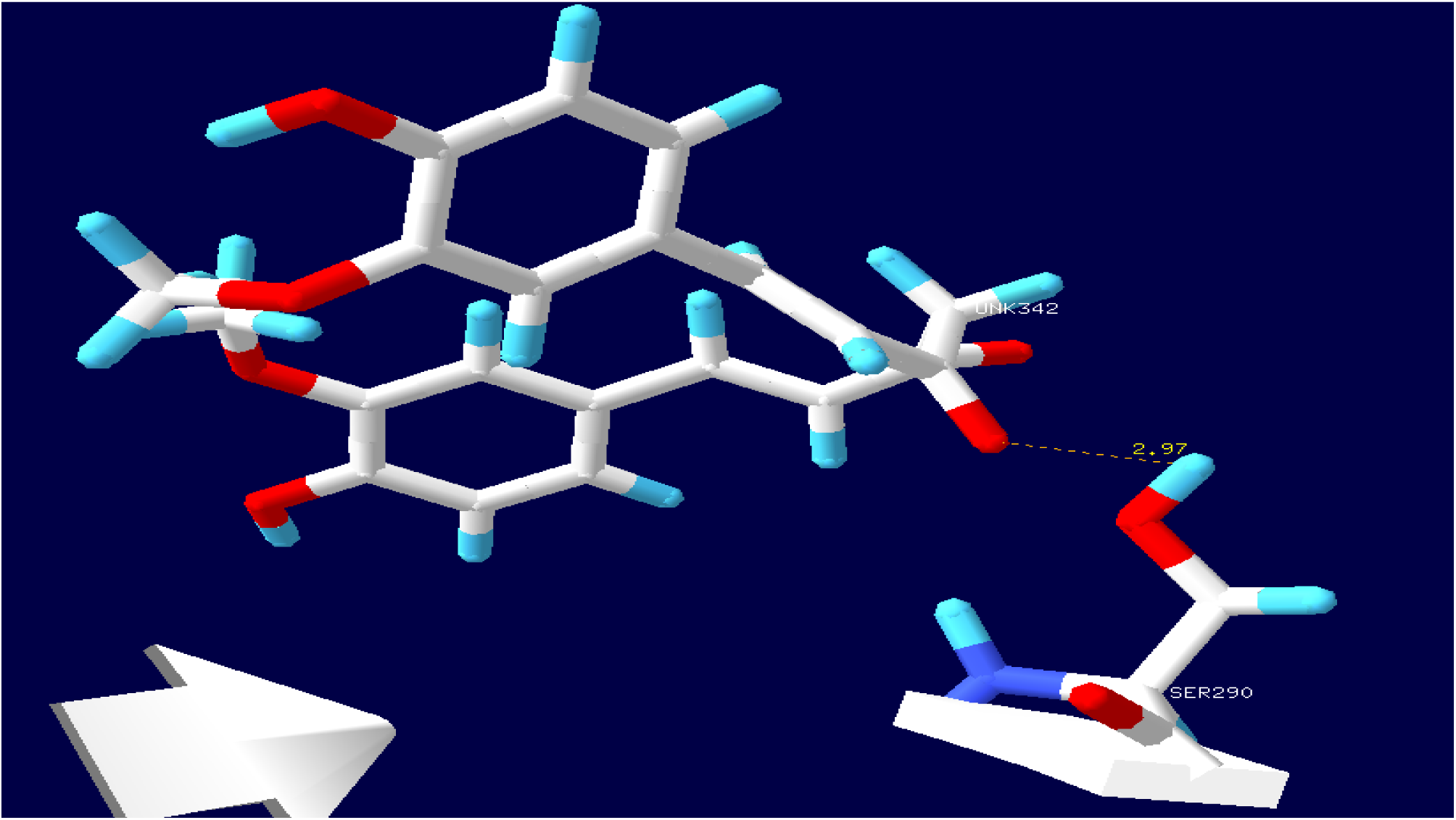
Curcumin in the active site of Nsp15 (5YVD A). *Swiss-PdbViewer* 4 1.0. Distance to Ser290 is 2.97 Å.

**Figure 15.**
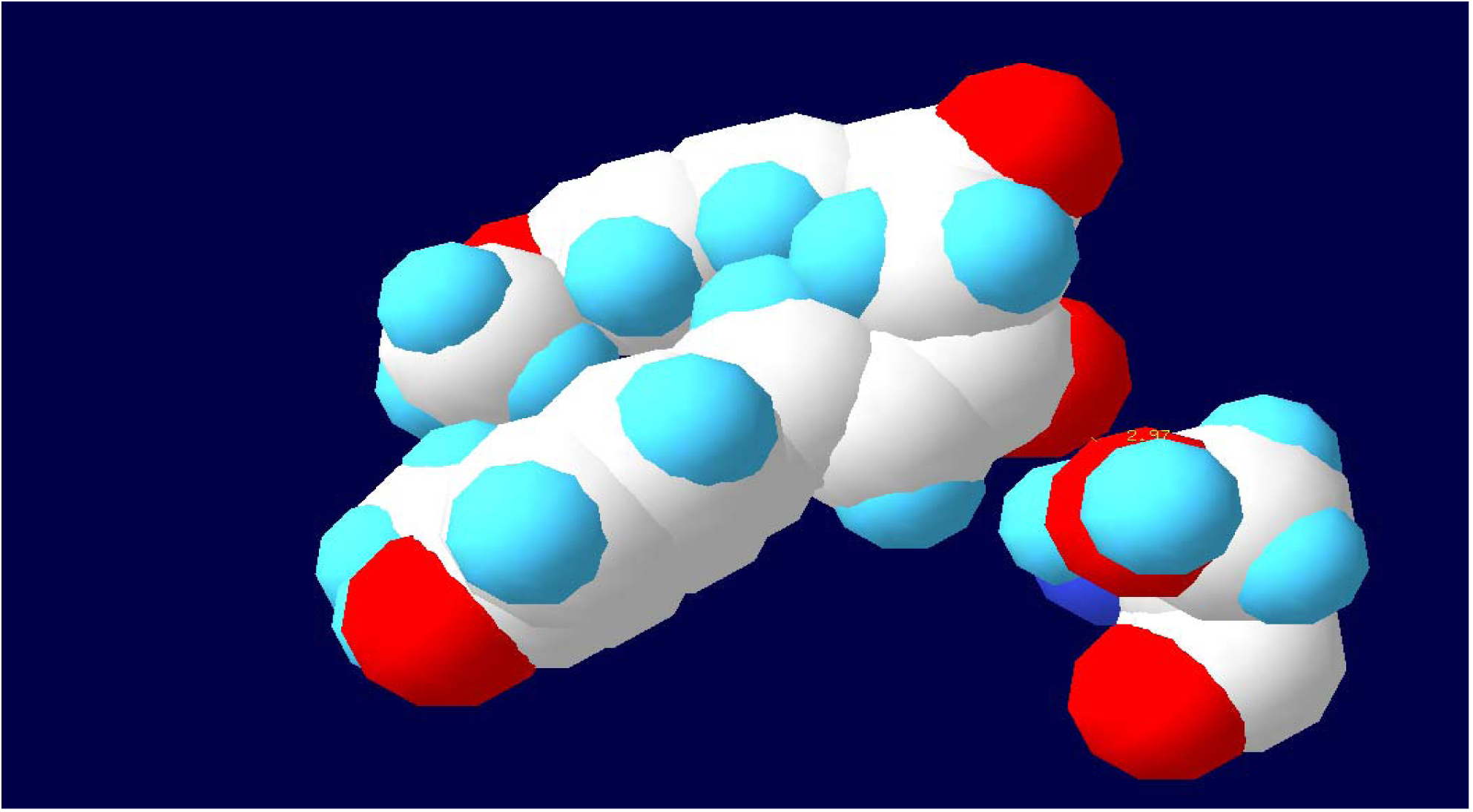
Curcumin in the active site of Nsp15 (5YVD A). *Swiss-PdbViewer* 4 1.0. Distance to Ser290 is 2.97 Å. Space filling model. On the left Curcumin, on the right bottom Ser290. This is to show proximity to Ser290.

**Figure 16.**
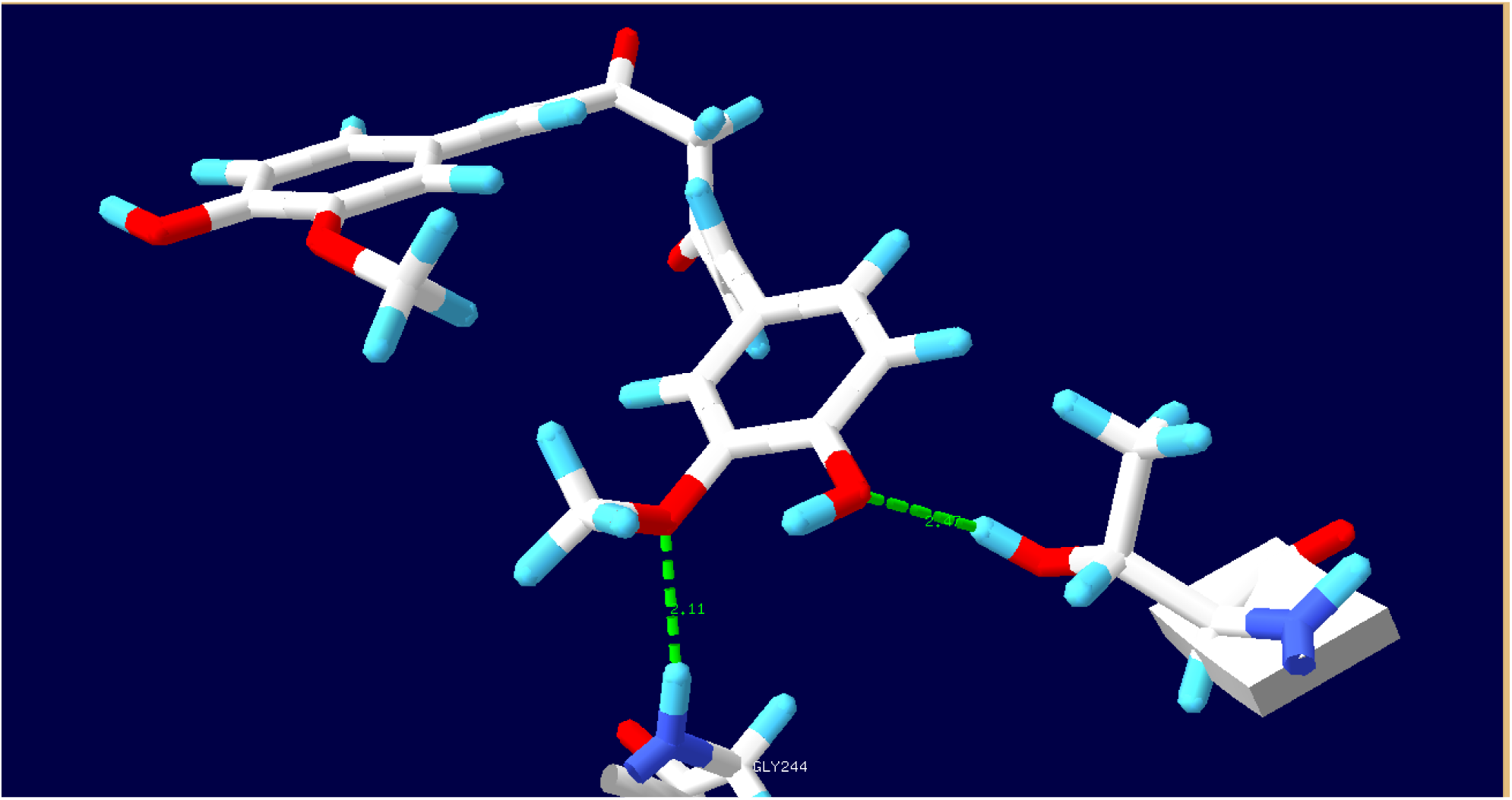
Curcumin in the active site of Nsp15 (5YVD A). *Swiss-PdbViewer* 4 1.0. Ligand hydrogen bond to Gly244 2.11 Å and Thr337 2.47 Å

**Figure 17.**
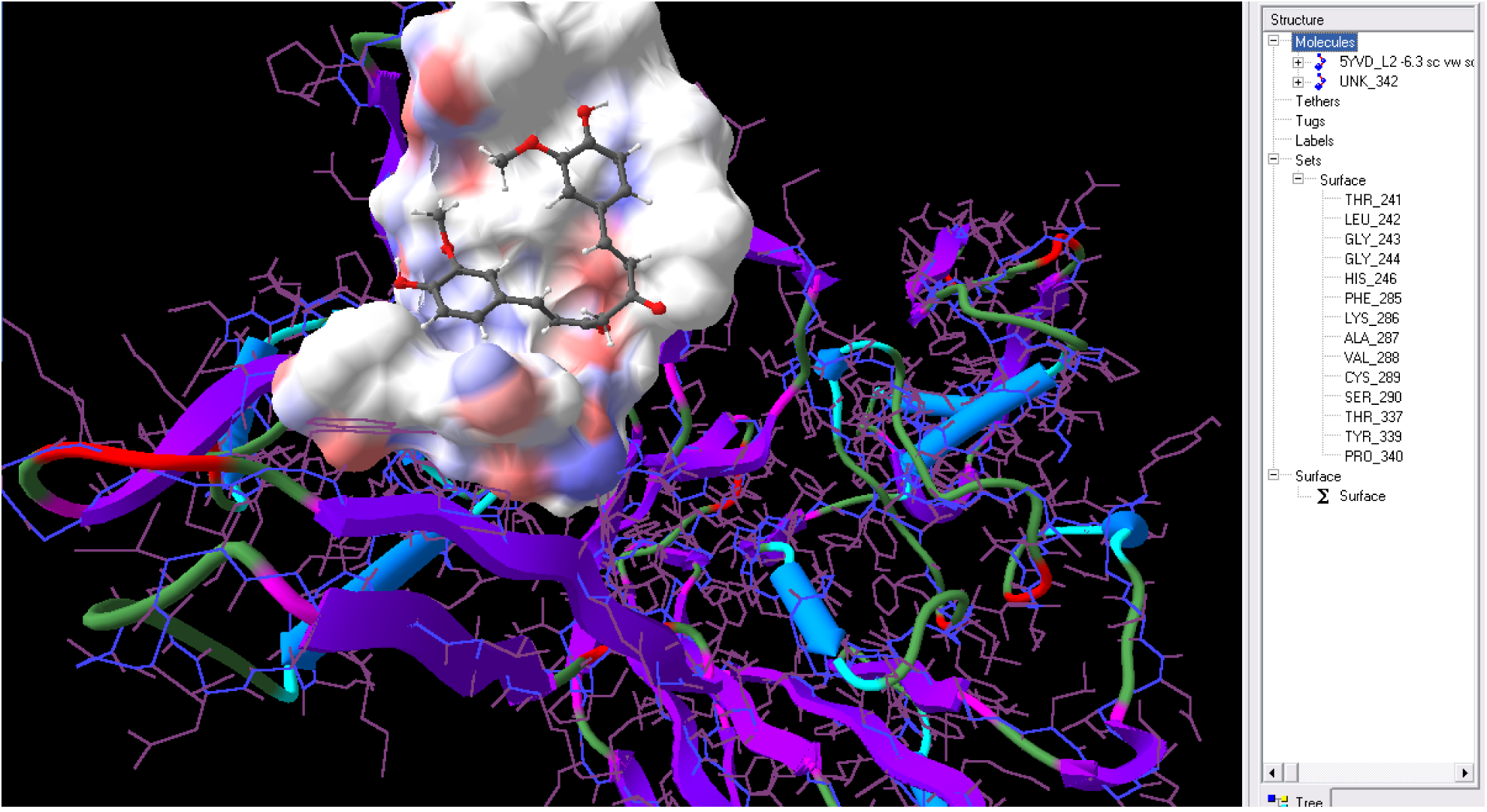
Curcumin in the active site of Nsp15 (5YVD A). *Sculpt* 3.0. Active site amino acids shown on the right.

**Figure 18.**
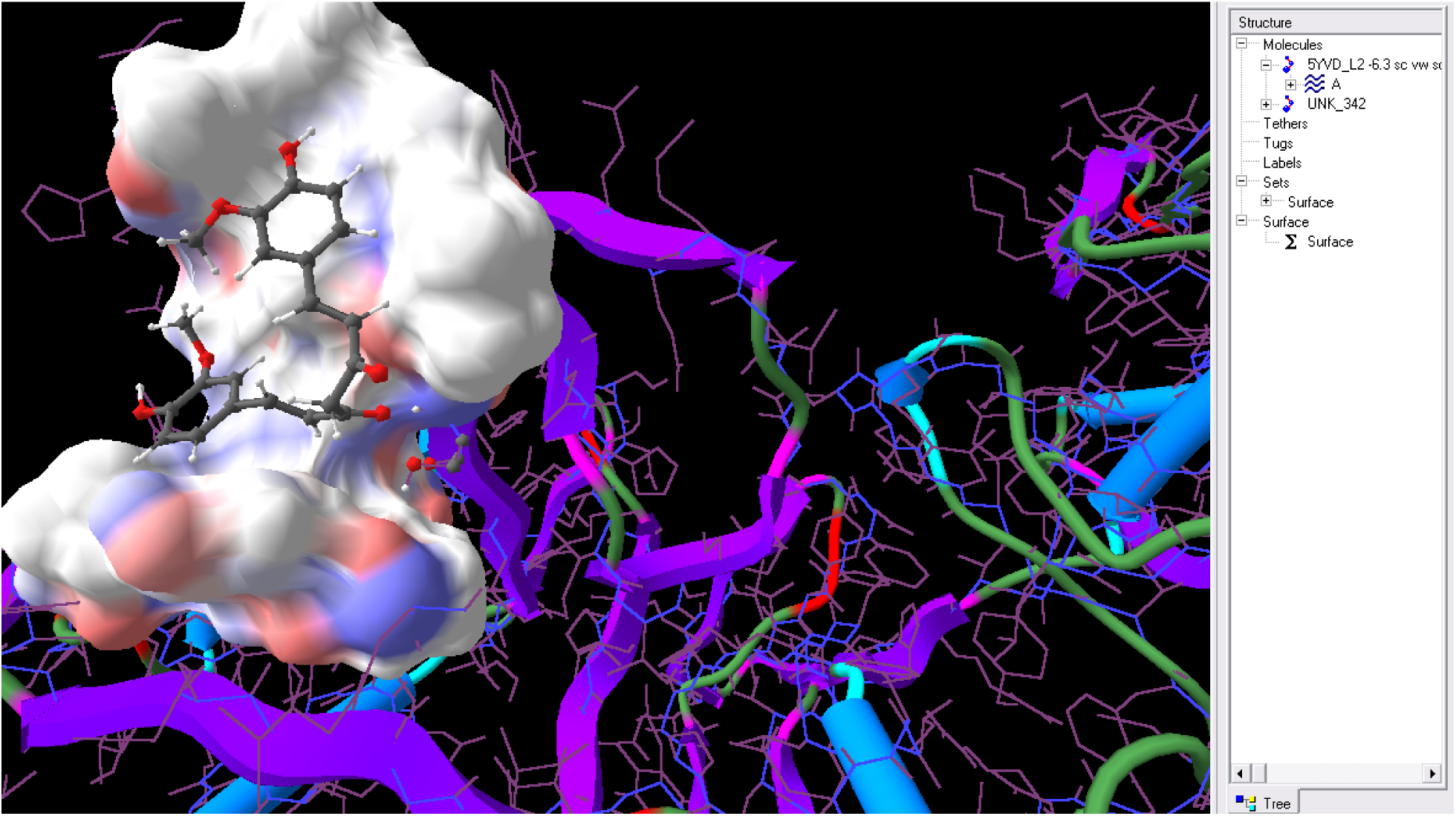
Curcumin in the active site of Nsp15 (5YVD A). Sculpt 3.0. Surface removed from Ser290 to show proximity.

### Results for Curcurcumin boronic acid derivative (CurcuminB)

**Figure 19.**
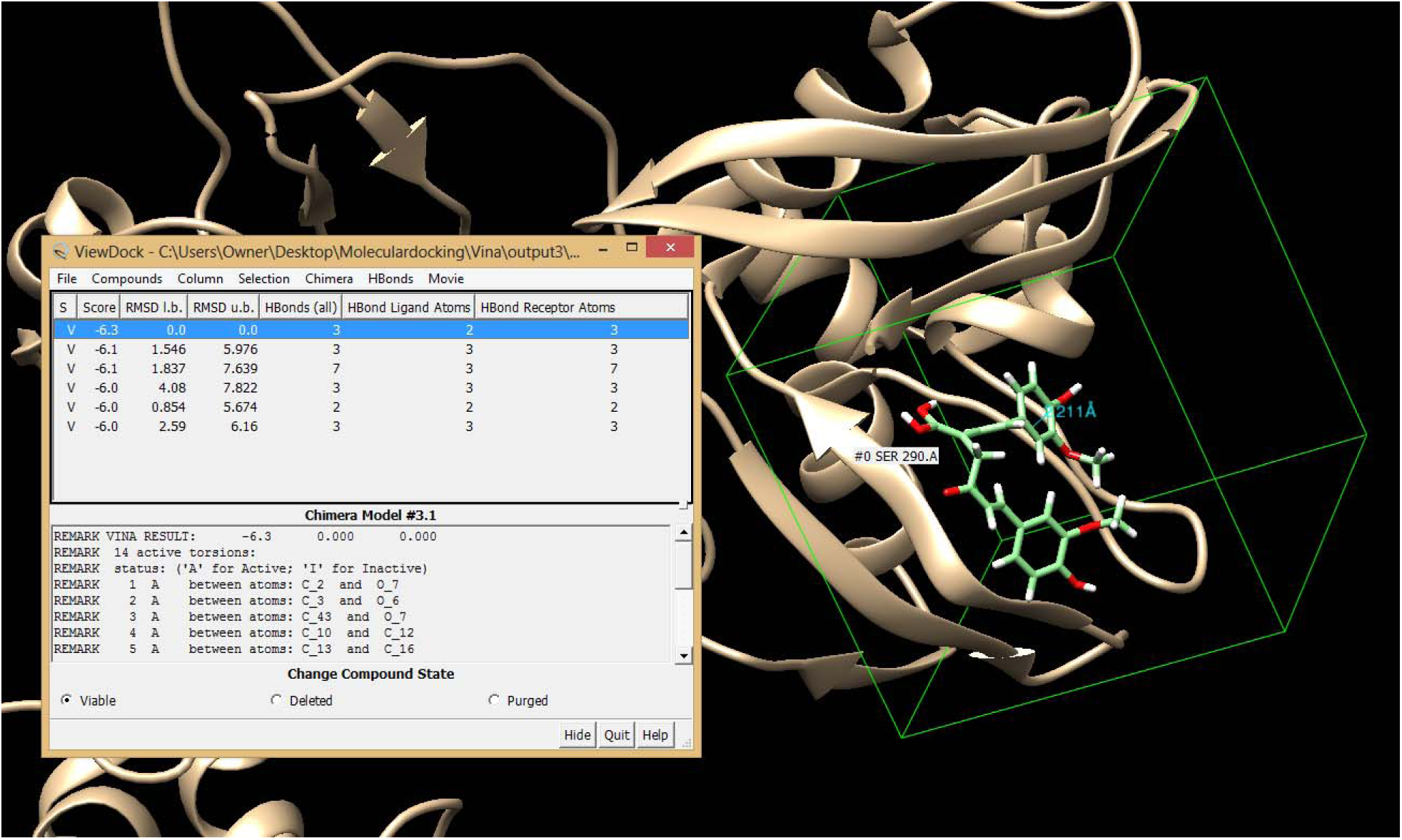
First take after CurcuminB was docked to 5YVD A with *AutoDock Vina. UCSF Chimera* 1.12

**Figure 20.**
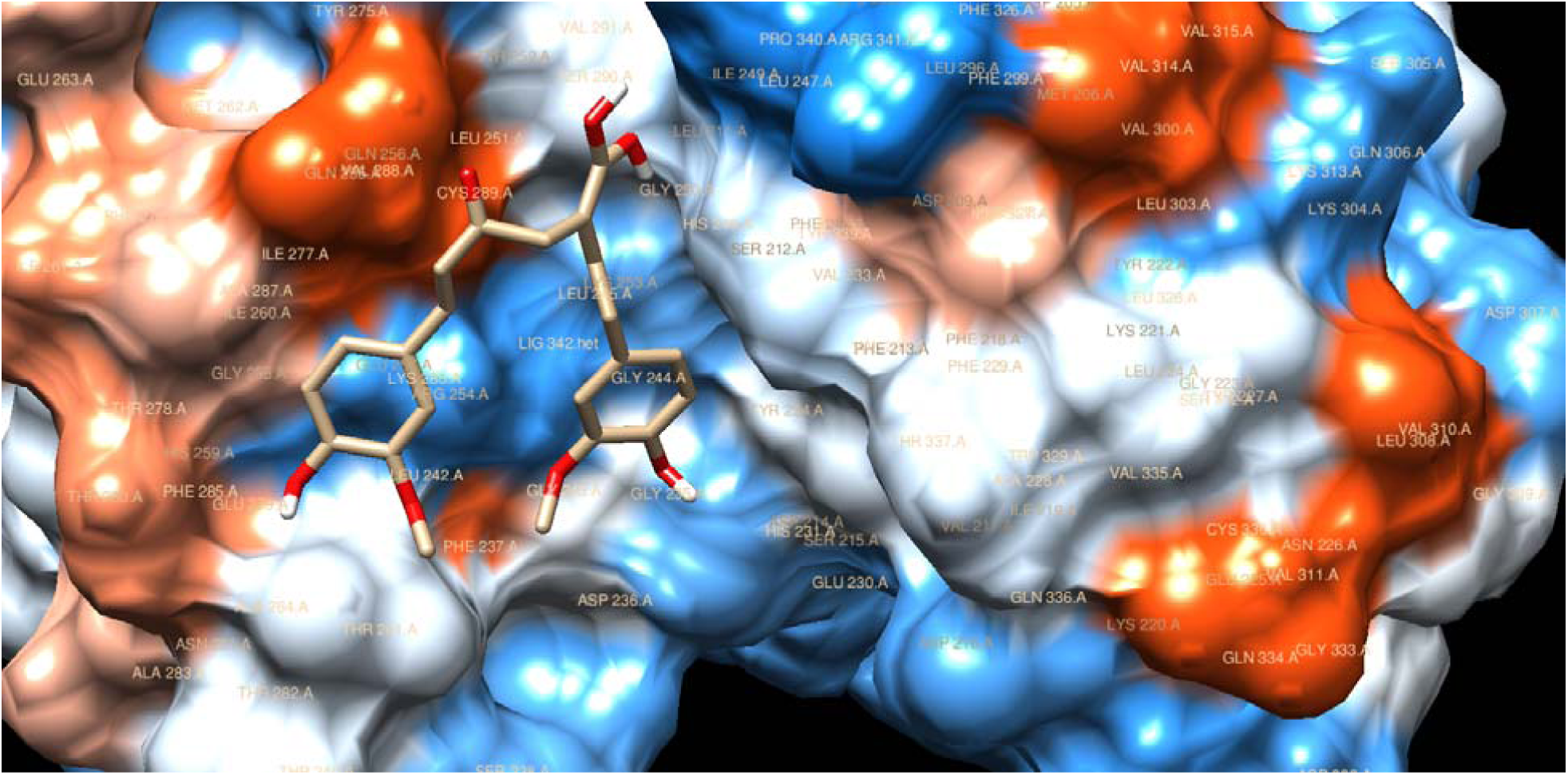
CurcuminB proximal to Ser290 in 5YVD A. *UCSF Chimera* 1.12.

**Figure 21.**
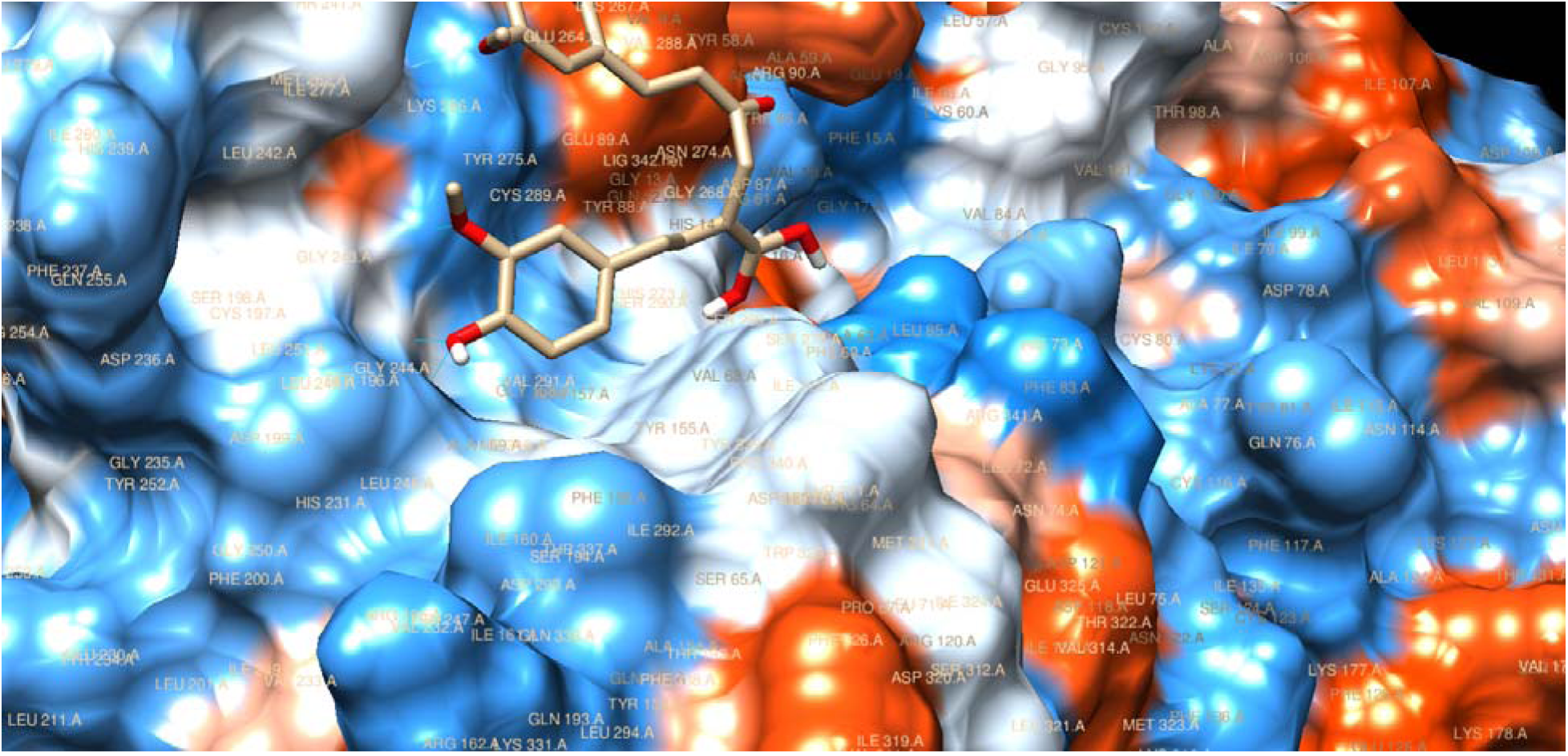
CurcuminB in proximity to a pocket in 5YVD A formed by His231 and Arg341 from which it forms two hydrogen bonds stabilizing the docking as if in an oxyanion hole. *UCSF Chimera* 1.12

**Figure 22.**
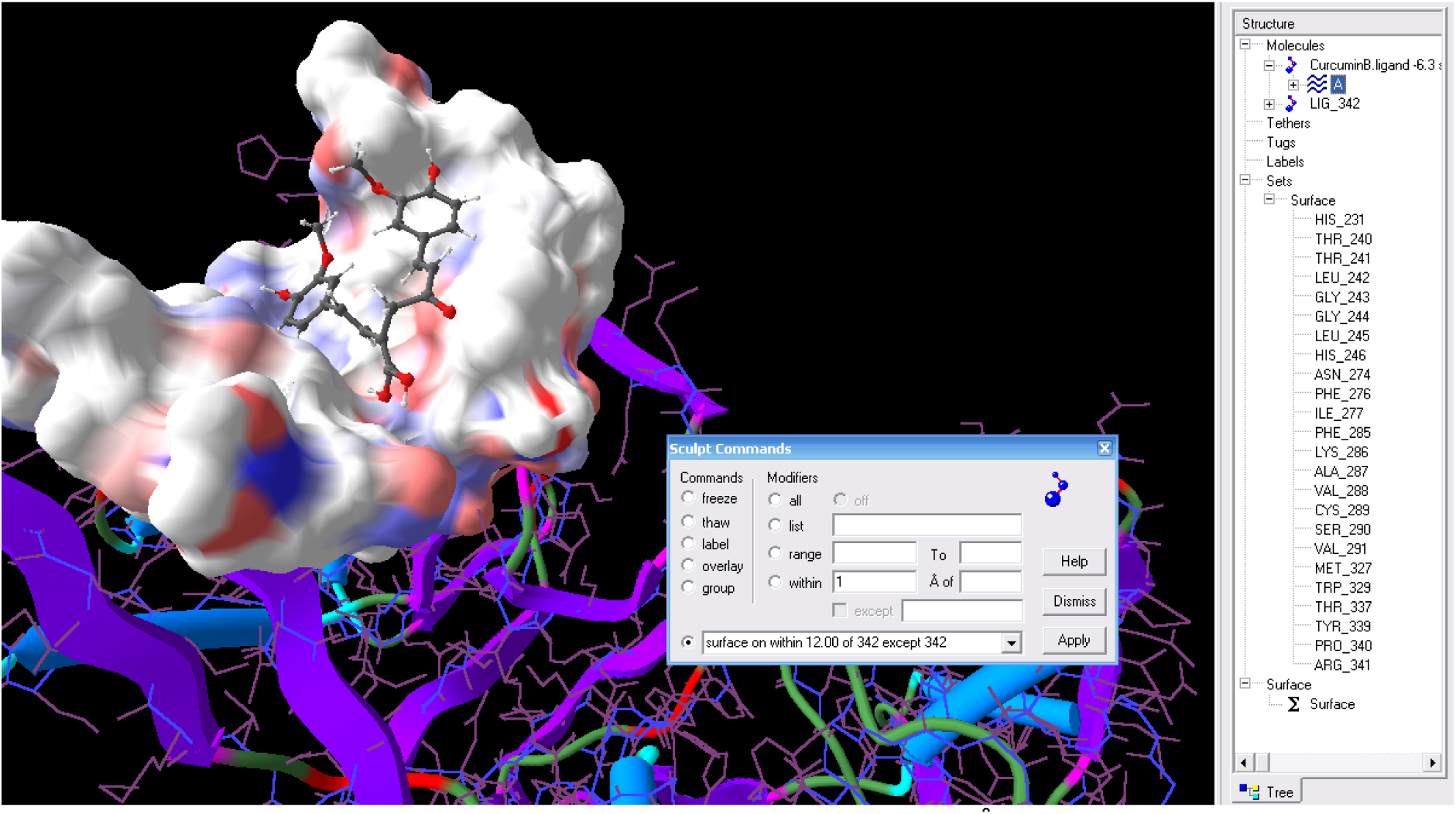
CurcuminB in the active site of 5YVD A. Surface on 12.00 Å from the ligan. This was to show His231 and Arg341 that appeared at 11.00 and 12.00 Å respectively. They form hydrogen bonds with the ligan. *Sculpt* 3.0.

**Figure 23.**
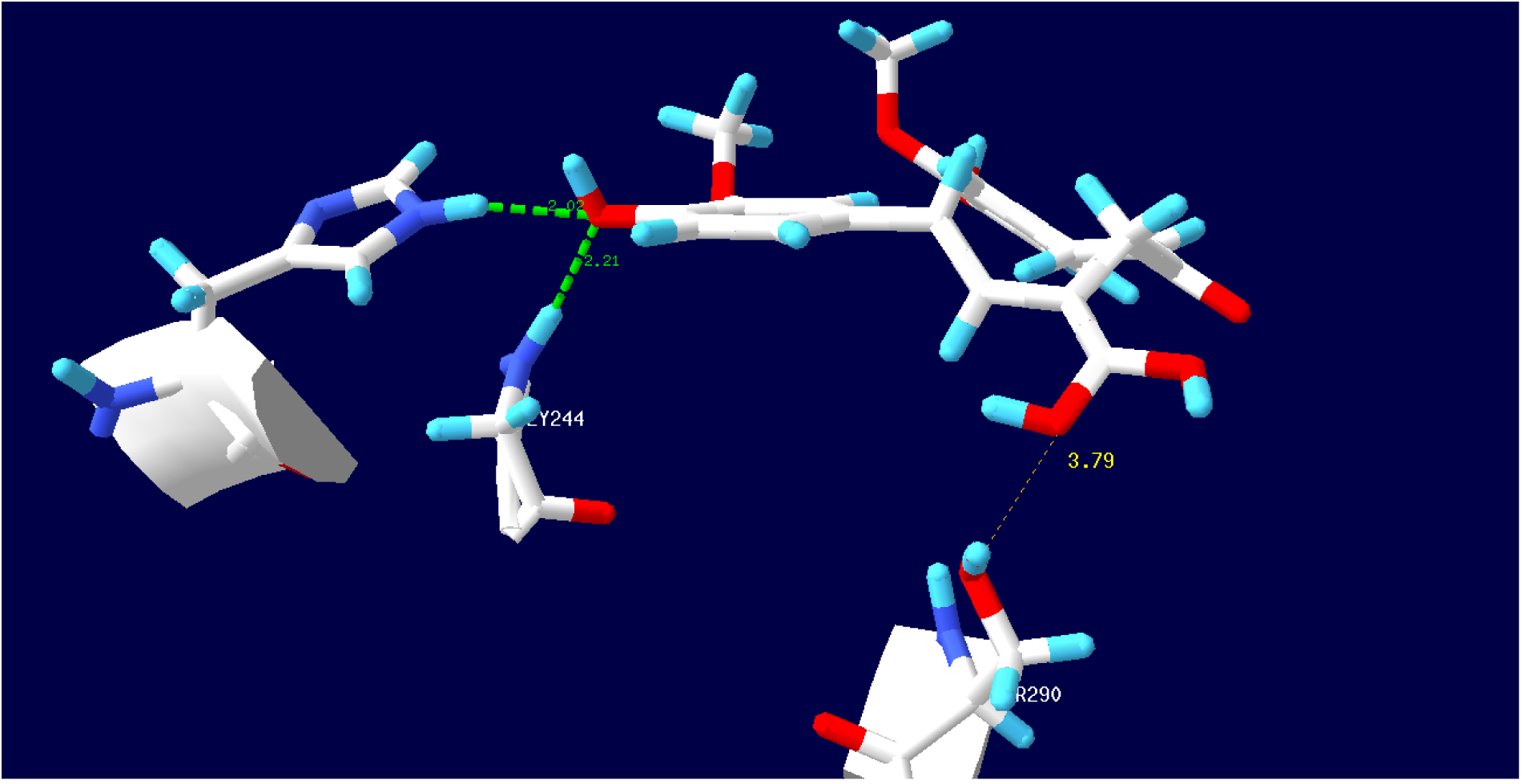
Distance from the Boron (Carbon in simulation) to the sidechain Oxygen in Ser90 of 5YVD A. Also showing the oxyanion like stabilization pocket formed between His231 and Gly244. *Swiss-PdbViewer* 4 1.0.

### Results for Hydroxychloroquine

**Figure 24.**
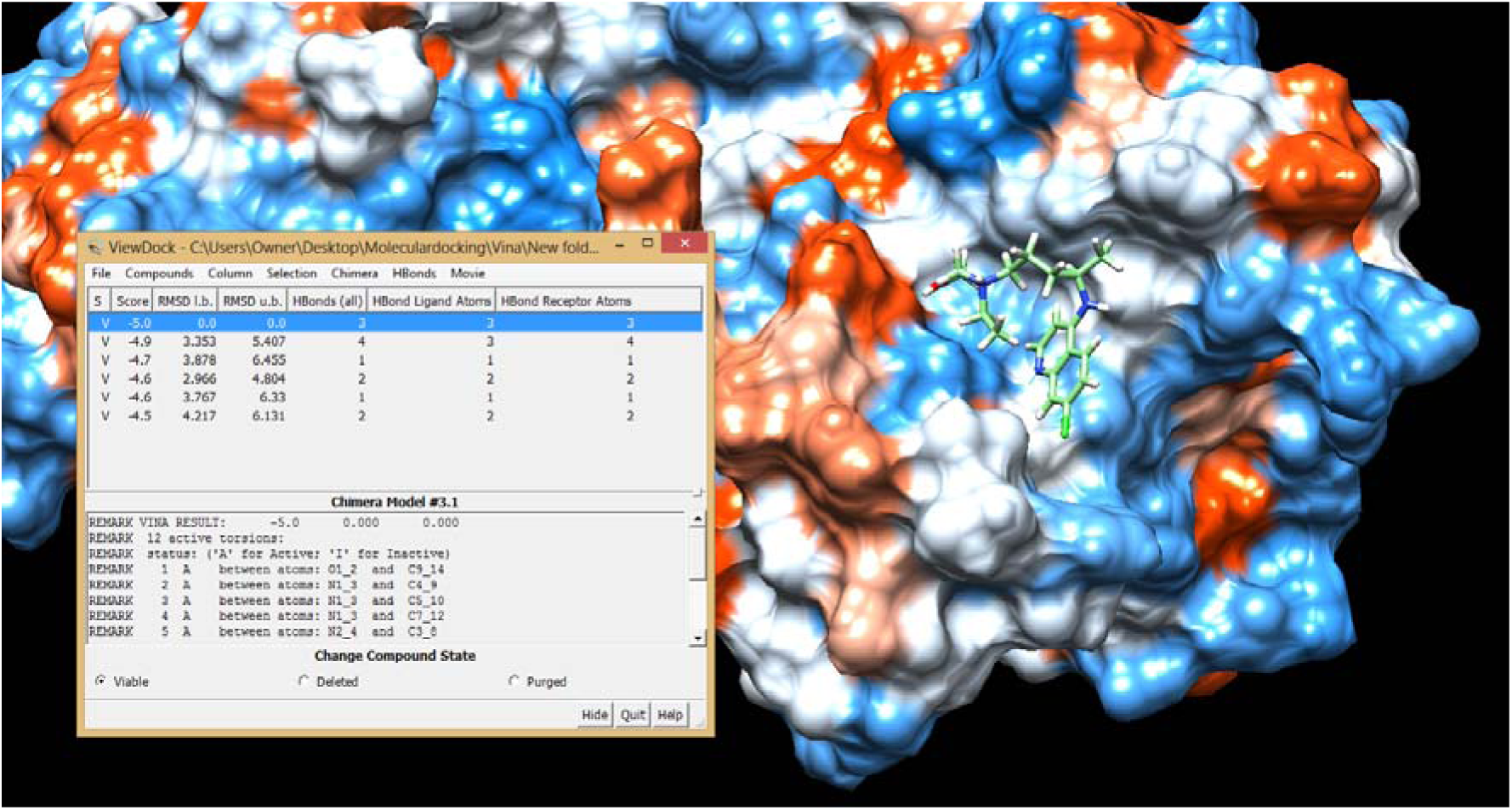
Hydroxychloroquine in the active site of 5YVD A. *UCSF Chimera* 1.12

### Results for HydroxychloroquineB

**Figure 25.**
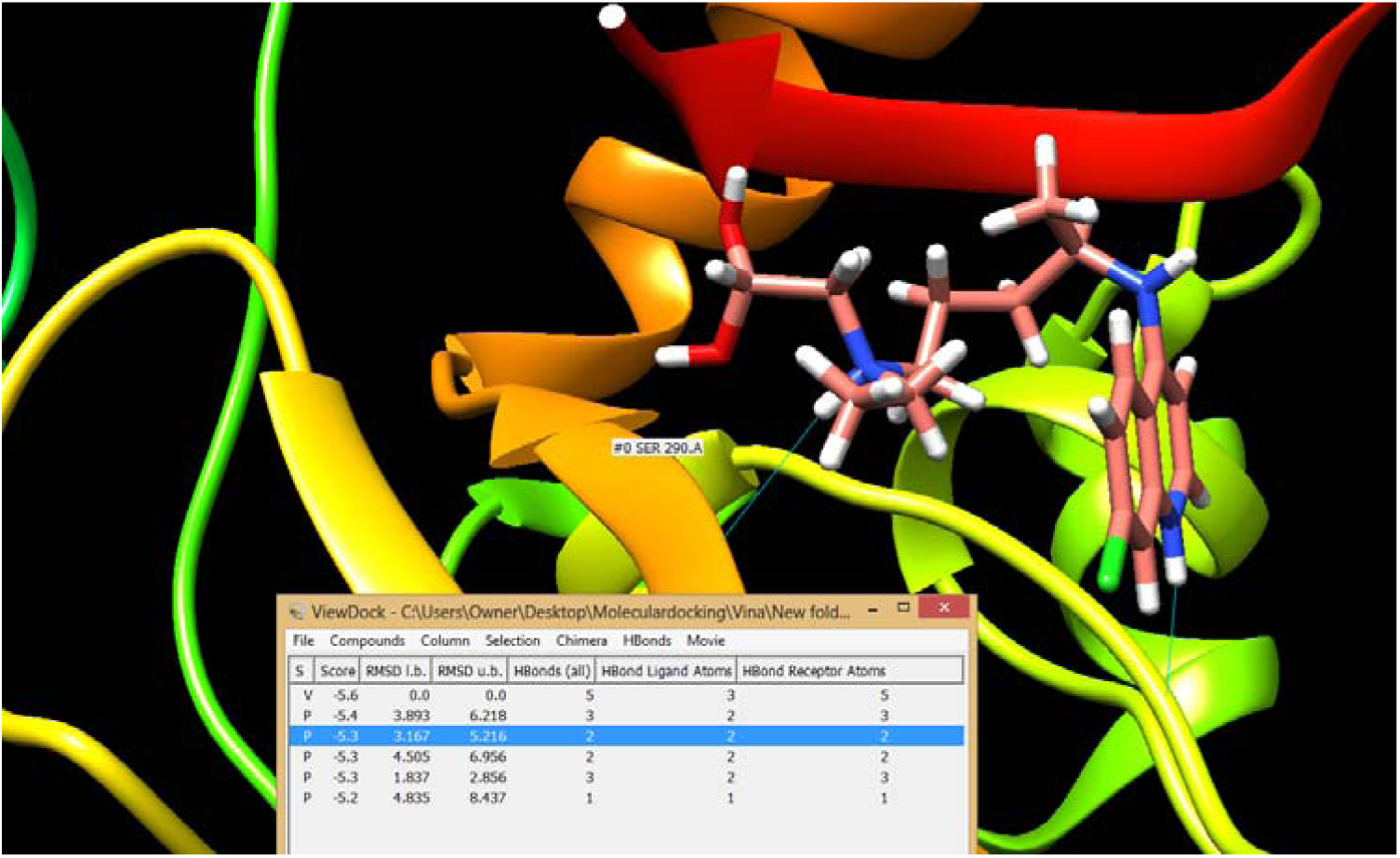
HydroxychloroquineB in the active site of 5YVD A. *USCF Chimera* 1.12

In this case the borono moiety best orients to Ser290 in the third outcome. Note that energy is −5.3 which is better than −5.0 of the Hydroxychloroquine best result.

**Figure 26.**
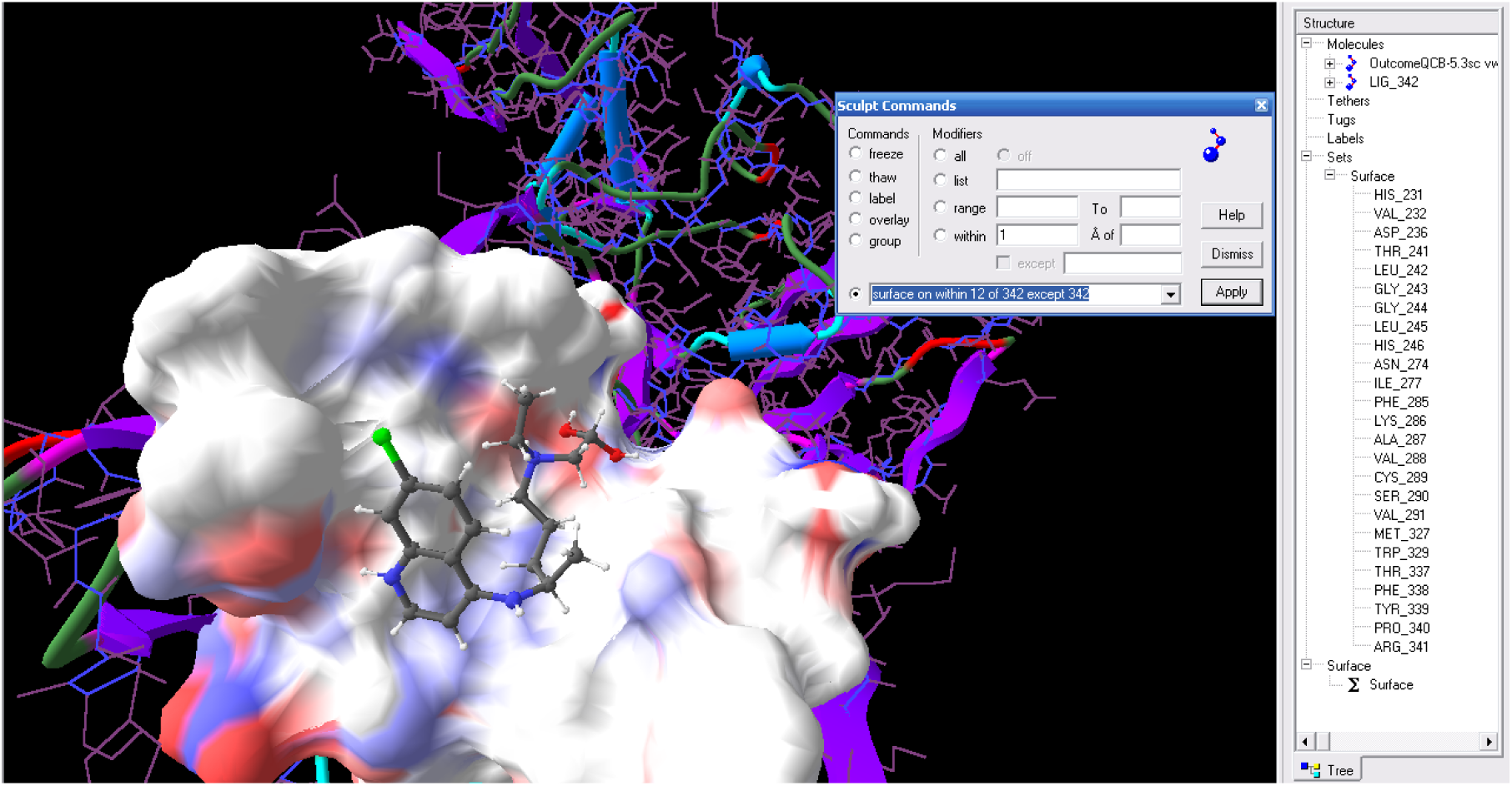
HydroxychloroquineB in the active site of 5YVD A. *Sculpt* 3.0. List of amino acids within 12.0 Angstroms. Note the presence of Ser290.

### Results for Glycerol

**Figure 27.**
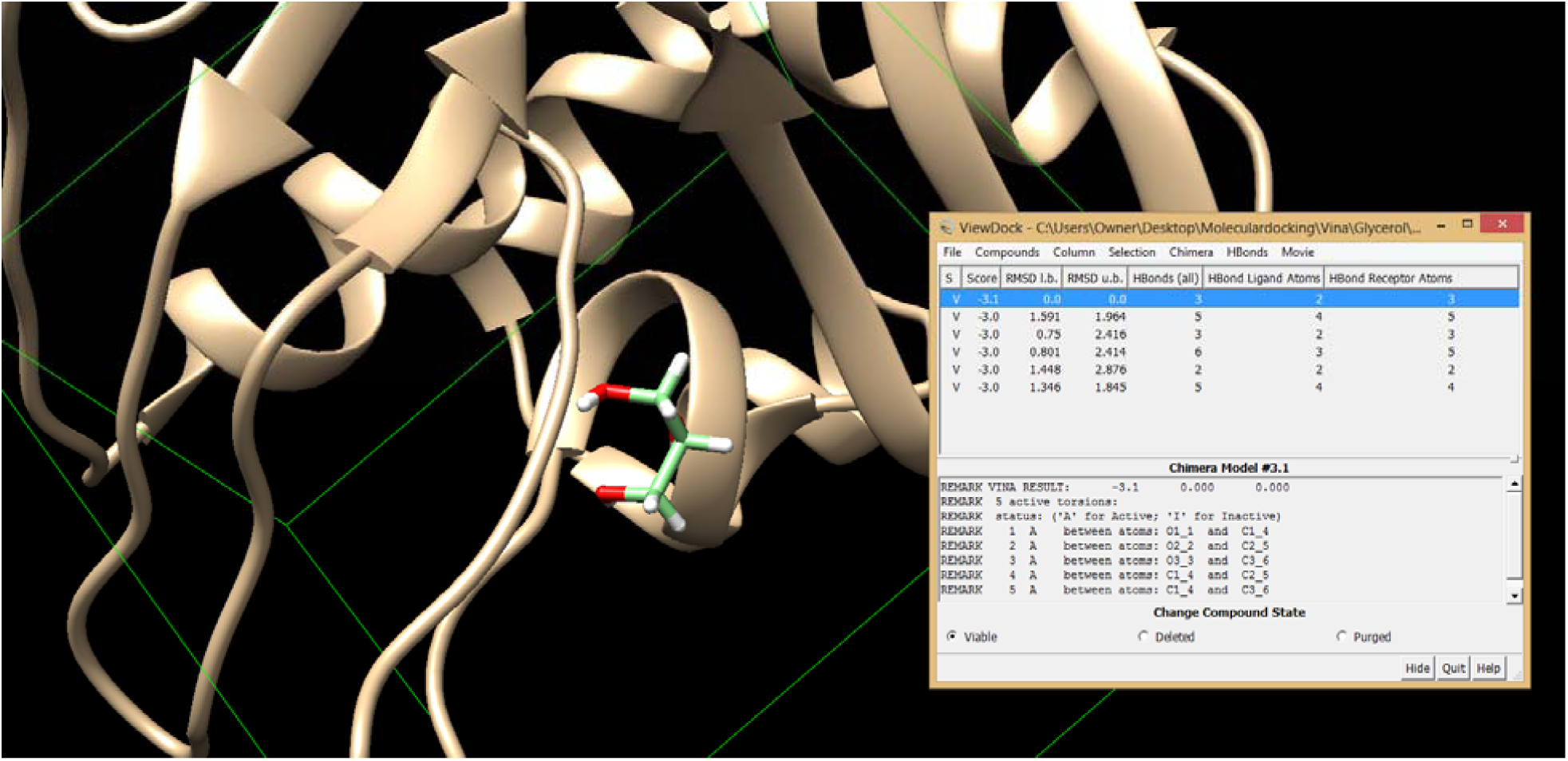
Glycerol in the active site of 5YVD A. *USCF Chimera* 1.12.

**Figure 28.**
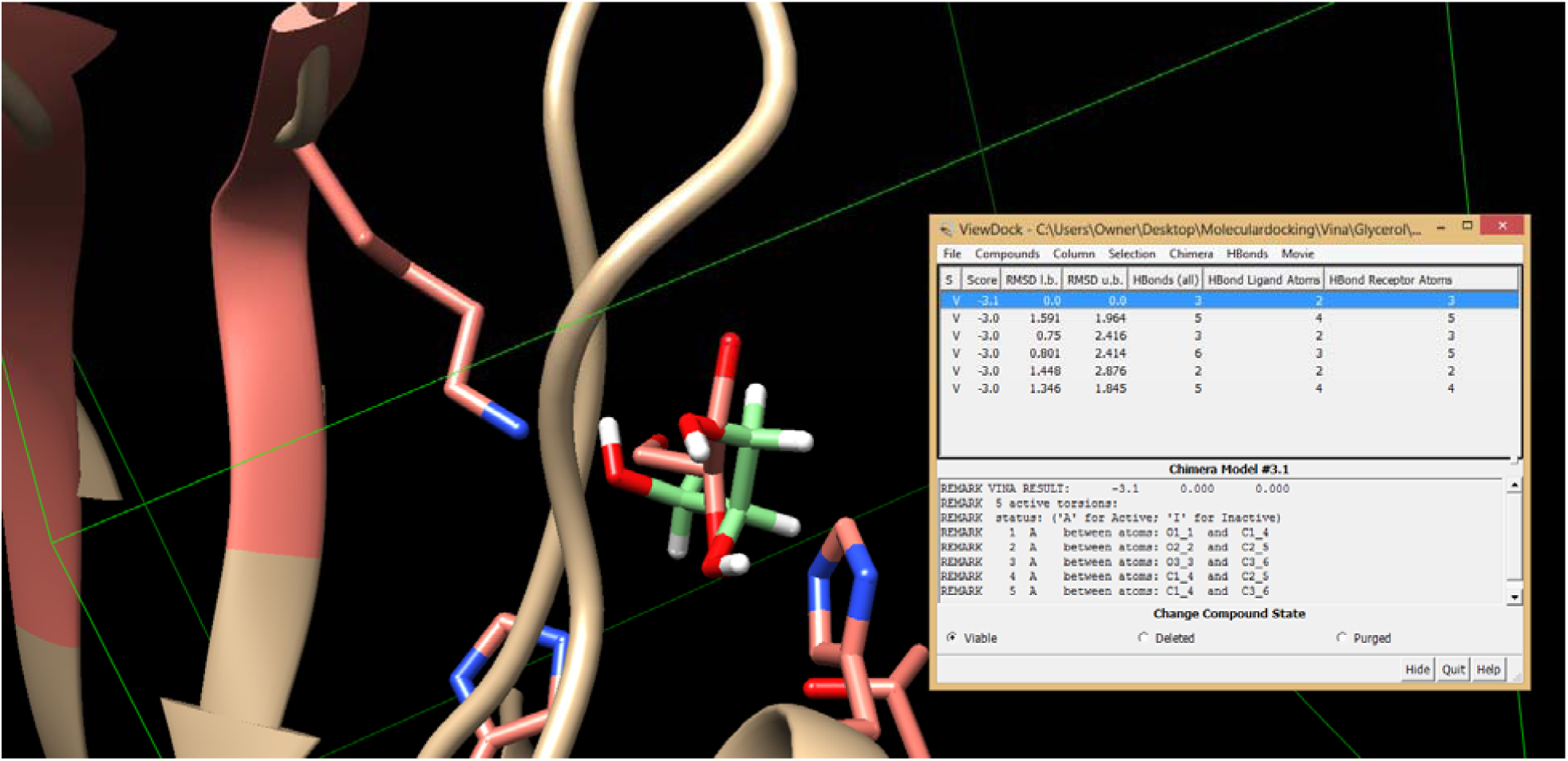
Glycerol in the active site of 5YVD A per our docking (green) vs. the NCBI structure (pink). This proves that *AutoDock Vina* is reliable for our purpose.

## Discussions and conclusions

1. Both compound, *p-*coumaric acid and curcumin that are found in nature fit within reasonable paramerters in the active site of Nsp15.
2. Both natural compounds dock with proximity to Ser290.
3. The enzyme may be treated as a serine protease in that uses serine 290 in the functions of endoribonuclease.
4. If we substitute one carbon atom or an oxygen, in the ligand, for a borono B(OH)_2_ moiety we may convert the compounds into boronic acids.
5. The boronic acids, being Lewis acids because of boron electron deficiency, will coordinate with Ser290. Therefore, showing improvement over their natural analogues.
6. In CurcuminB, there is greater stabilization by the two hyddrogen bonds from Gly244 (2.211 Å), and His231 (2.017 Å) to the ligand acting as some sort of oxyanion hole. Therefore, CurcuminB should be consider a highly probable potent inhibitor of Nsp15.
7. Both Hydroxychloroquine and HydroxychloroquineB fit within reasonable parameters in the active site of this enzyme. However, they are both less effective than our best compounds Curcumin and CurcuminB per *AutoDock Vina*’s results.

## Aknowledgment

My special gratitude to my former mentor Prof. Manfred Philipp, Ph.D. retired professor from Lehman College and the Graduate Center/CUNY for his support in my endeavor.

## Appendix A.

### Stuctures downloaded from internet

**Figure A1.**
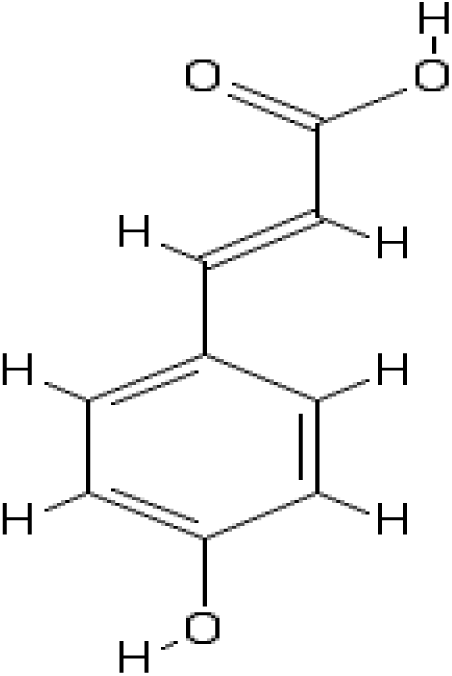
*p*-coumeric acid, (*PubChem*). ID: 637542

**Figure A2.**
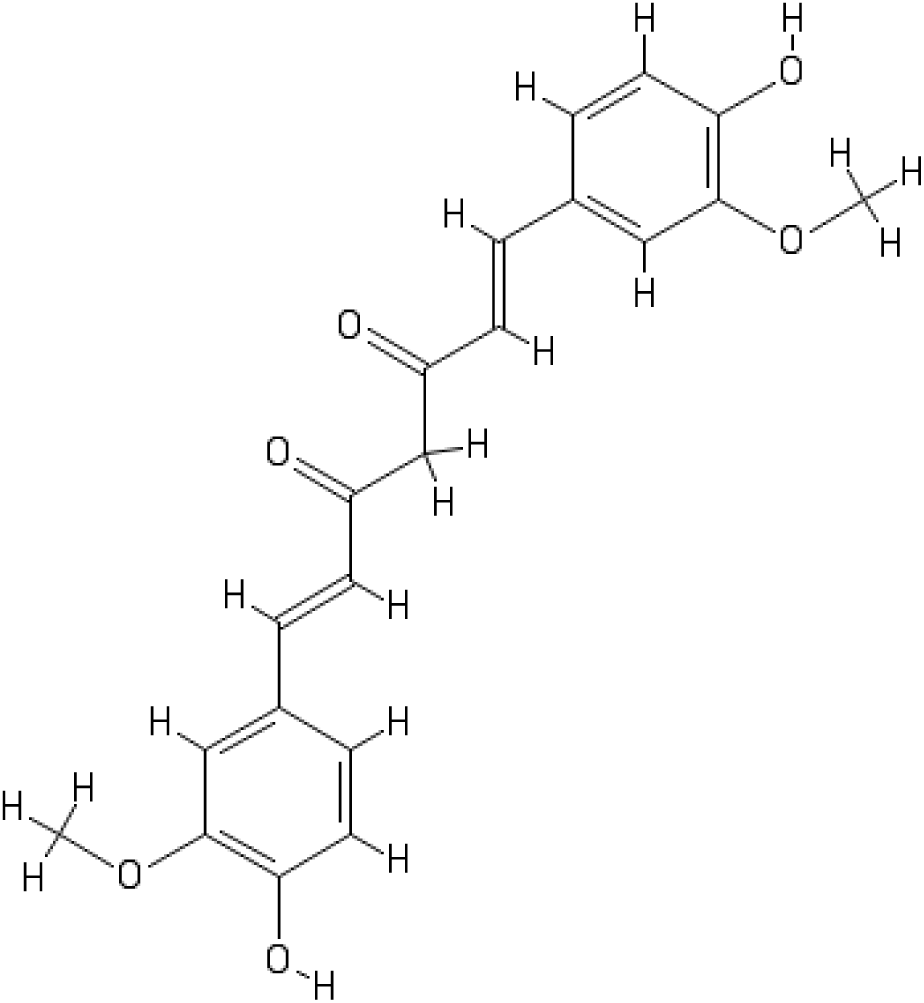
Curcumin. PubChem from NCBI. CID: 969516 (*PubChem*).

